# An optimized Tet-On system for conditional control of gene expression in sea urchins

**DOI:** 10.1101/2022.10.12.511941

**Authors:** Jian Ming Khor, Charles A. Ettensohn

## Abstract

Sea urchins and other echinoderms are important experimental models for studying developmental processes. The lack of approaches for conditional gene perturbation, however, has made it challenging to investigate the late developmental functions of genes that have essential roles during early embryogenesis and genes that have diverse functions in multiple tissues. The doxycycline-controlled Tet-On system is a widely used molecular tool for temporally and spatially regulated transgene expression. Here, we optimized the Tet-On system to conditionally induce gene expression in sea urchin embryos. Using this approach, we explored the roles the MAPK signaling plays in skeletogenesis by expressing genes that perturb the pathway specifically in primary mesenchyme cells (PMCs) during later stages of development. We demonstrated the wide utility of the Tet-On system by applying it to a second sea urchin species and in cell types other than the PMCs. Our work provides a robust and flexible platform for the spatio-temporal regulation of gene expression in sea urchins, which will considerably enhance the utility of this prominent model system.

## Introduction

Sea urchin embryos are a prominent model system for investigating developmental processes, due to their relatively simple organization, optical transparency, and amenability to experimental manipulation. During development, diverse cell types with transient regulatory states are defined by networks of differentially expressed genes, structured as modular gene regulatory networks or GRNs. The core components of GRNs are comprised of temporally and spatially restricted transcription factors (TFs), signaling factors, and the downstream terminal differentiation genes controlled by the regulatory genes. Determining how these combinations of genes, operating within a hierarchical network, drive the morphogenesis of specific anatomical features during development will lead to a better understanding of the connection between genotype and phenotype.

To establish the causal links required for the construction of experimental GRN models, functional perturbations of regulatory genes are usually performed, followed by observation of the effects on potential target genes. Although several approaches have been employed to successfully perturb gene function in sea urchin embryos, functional studies generally rely on microinjection of reagents such as morpholino antisense oligonucleotides (MOs) to block translation or splicing (Yaguchi, 2019), mRNAs encoding dominant-negative or constitutively active forms of proteins (Lepage and Gache, 2004), and targeted genome editing nucleases, such as transcription activator-like effector nucleases (TALENs) (Yamazaki et al., 2021) and CRISPR-Cas9 (Fleming et al., 2021; Lin et al., 2019; Wessel et al., 2020). These conventional approaches, however, face a common constraint. Most notably, they do not allow for conditional gene perturbation. As microinjection is normally performed on fertilized eggs, the injected reagent will exert its effects ubiquitously and almost immediately, which consequently limits this approach to early developmental stages. Chemical inhibitors provide more flexibility in the timing of their application, but may lack molecular specificity or produce on-target side effects as they commonly target signaling pathways that regulate diverse cellular functions such as cell growth, proliferation, and differentiation. A modified version of MOs that can cross cell membranes, termed Vivo-MOs, have been reported to effectively silence genes in sea urchin embryos (Heyland et al., 2014; Luo and Su, 2012), but their use has been limited by low solubility and toxicity (Cui et al., 2017). More recently, caged morpholinos have been used for temporally regulated gene knockdowns in sea urchins (Bardhan et al., 2021).

The skeletogenic GRN that drives the formation of the intricate, calcite-based endoskeleton in sea urchin larvae has been intensively investigated (see reviews by McIntyre et al., 2014; Shashikant et al., 2018). The skeleton is produced by primary mesenchyme cells (PMCs), which are descendants of the four large micromeres that form during early development through the actions of asymmetric division and localized maternal factors. At the mesenchyme blastula stage, the PMCs undergo epithelial-mesenchymal transition (EMT) and ingress into the blastocoel. As the embryo begins to gastrulate, a cell-autonomous program that initially deploys the skeletogenic GRN gradually shifts to become signal-dependent. During gastrulation, ectoderm-derived cues, such as vascular endothelial growth factor-3 (VEGF3), guide PMC migration and fusion within the blastocoel to form a distinctive ring-like pattern consisting of two clusters of cells (the ventrolateral clusters) (Adomako-Ankomah and Ettensohn, 2013; Duloquin et al., 2007; Knapp et al., 2012; Morgulis et al., 2019; Sun and Ettensohn, 2014). It is within these two cell clusters that spicule formation is initiated. During later stages of development, signals from the adjacent ectodermal cells continue to stimulate the growth, elongation, and branching of skeletal rods that extend from the spicule primordia. Treatment with axitinib, a selective VEGF receptor inhibitor, revealed that VEGF signaling is required for early ingression and also PMC migration and patterning during later stages of development (Adomako-Ankomah and Ettensohn, 2013; Morgulis et al., 2019; Morgulis et al., 2021). Genes with localized expression at the tips of the growing arms are also downregulated in axitinib-treated embryos (Morgulis et al., 2019; Sun and Ettensohn, 2014; Tarsis et al., 2022).

The mitogen-activated protein kinase (MAPK) signaling pathway has been shown to be required for sea urchin embryonic skeletogenesis (Fernandez-Serra et al., 2004; Röttinger et al., 2004). Phosphorylated extracellular signal-regulated kinase (ERK), a marker for MAPK pathway activity, is found in the micromere lineage prior to PMC ingression during early development. During gastrulation, activated ERK is detected in the PMC ventrolateral clusters and in the adjacent ectoderm, as well as in secondary mesenchyme cells (SMCs) at the tip of the invaginating archenteron. During later stages of larval development, ERK activation is observed in diverse cell types such as the coelomic pouches and foregut (Fernandez-Serra et al., 2004). Inhibition of mitogen-activated protein kinase kinase (MEK), an upstream activator of ERK, using U0126 blocks PMC ingression and differentiation (Fernandez-Serra et al., 2004; Röttinger et al., 2004). Overexpression of either dominant negative MEK or dual specificity phosphatase 6 (DUSP6, also known as MAP kinase phosphatase 3 or MKP3), both of which are known to downregulate ERK activity, also results in inhibition of PMC specification (Fernandez-Serra et al., 2004; Röttinger et al., 2004).

In previous studies, activation of the MAPK signaling pathway in the large micromere lineage was shown to be cell autonomous. In the early embryo, MAPK signaling is regulated by maternal β-catenin (Röttinger et al., 2004). Additionally, U0126 treatment blocks the formation of spicules and downregulates the expression of biomineralization genes in micromere cultures (Fernandez-Serra et al., 2004). Activated ERK is detected in dissociated blastomeres prior to the hatched blastula stage, at roughly the same time as in control embryos (Röttinger et al., 2004). During later stages of development, dissociated blastomeres exhibit lower levels of activated ERK compared to intact embryos, suggesting that other regulatory inputs are necessary for sustained MAPK activation following the shift from cell-autonomous to signal-dependent regulation of development. In the same studies, skeletogenesis was shown to be inhibited in embryos exposed to U0126 after PMC ingression. Embryos treated with U0126 during later stages of embryonic development exhibit downregulation of biomineralization genes at the growing tips of the anterolateral and postoral rods (Sun and Ettensohn, 2014). As many studies in mammalian systems have revealed that VEGF induces MAPK signaling to stimulate various cellular functions (Doanes et al., 1999), these findings point to a possible link between VEGF and MAPK signaling during late stages of sea urchin skeletogenesis, although a direct connection has not been established.

Ets1, a key transcription factor in the sea urchin skeletogenic GRN, was identified as a putative target of ERK phosphorylation (Röttinger et al., 2004). RNA-seq studies have shown that many PMC effector genes are positively regulated by Ets1 (Rafiq et al., 2014). Ets1 expression also coincides with activated ERK during PMC and SMC ingression (Fernandez-Serra et al., 2004; Röttinger et al., 2004). Knockdown of Ets1 with Ets1 MO (Rafiq et al., 2012) or suppression of endogenous Ets1 activity through overexpression of a dominant negative form of Ets1 (Kurokawa et al., 1999; Sharma and Ettensohn, 2010), phenocopies the effects of U0126 treatment. In contrast, overexpression of a constitutively active form of Ets1 (phosphomimetic Ets1) restores PMC specification in embryos treated with U0126, suggesting that MAPK signaling regulates skeletogenesis through Ets1 phosphorylation (Röttinger et al., 2004).

Like many regulatory and signaling genes, ERK and Ets1 are expressed in diverse embryonic temporal and spatial domains (Fernandez-Serra et al., 2004; Röttinger et al., 2004). Hence, the lack of conditional approaches in the sea urchin model system has made it challenging to precisely define the developmental functions of these proteins. In the current study, we modified and optimized a two-plasmid Tet-On system for inducible transgene expression to explore the roles the MAPK signaling pathway plays in PMC specification and skeletogenesis. The tetracycline-responsive Tet-On and Tet-Off gene expression systems are widely used in many eukaryotic models to control transgene activity (see review by Das et al., 2016). The Tet-On system permits activation of transgene expression by treatment with tetracycline or tetracycline-derivatives like doxycycline (Dox), whereas the Tet-Off system permits transgene silencing through continuous Dox administration. The Tet-On system we used in this study consist of two different components: (1) a third-generation *reverse tetracycline-controlled transactivator* (rtTA or tetON3G) gene downstream of sea urchin-specific CRE and promoter and (2) the gene of interest downstream of the tetracycline response element (TRE) and the minimal CMV promoter. Technical advancements in inducible gene perturbation such as the work we presented here will considerably enhance the utility of sea urchins and other echinoderms as models for developmental studies.

## Results

### A two-plasmid Tet-On system confers tight control of transgene expression

The ability to regulate the timing of transgene expression in a cell type-specific manner is a powerful tool in developmental studies. To direct transgene expression spatially and temporally, we optimized a two-plasmid Tet-On system, consisting of a pair of transactivator and responder constructs (see Materials and Methods). We first placed the *reverse tetracycline-controlled transactivator* (*rtTA*) gene under the control of a *Sp-EMI/TM* intronic *cis*-regulatory element (CRE) (characterized in Khor et al., 2019) and the *Sp-endo16* ubiquitous promoter to generate the transactivator construct, PMC-CRE: rtTA (**Figure 1A**). We then placed the GFP coding sequence immediately downstream of the tetracycline response element (TRE) and the minimal CMV promoter to generate the responder construct TRE: GFP. As a preliminary test of the functionality of the Tet-On system, we co-injected the plasmids and exposed transgenic *L. variegatus* embryos to 5 μg/mL doxycycline (Dox) (**Figure S1A**). Transgenic embryos exposed to Dox overnight showed strong GFP fluorescence exclusively in the PMCs (**Figure 1B**). The distribution of GFP matched that observed when GFP is under direct, constitutive control of the *Sp-EMI/TM* CRE (Khor et al., 2019). As GFP protein can readily diffuse throughout the PMC syncytium, the entire PMC network is labeled in transgenic embryos, despite the mosaic incorporation and expression of transgenes in sea urchins. In the absence of Dox, GFP protein expression was undetectable, indicating that our Tet-On system conferred tight control of gene expression.

**Figure 1:**
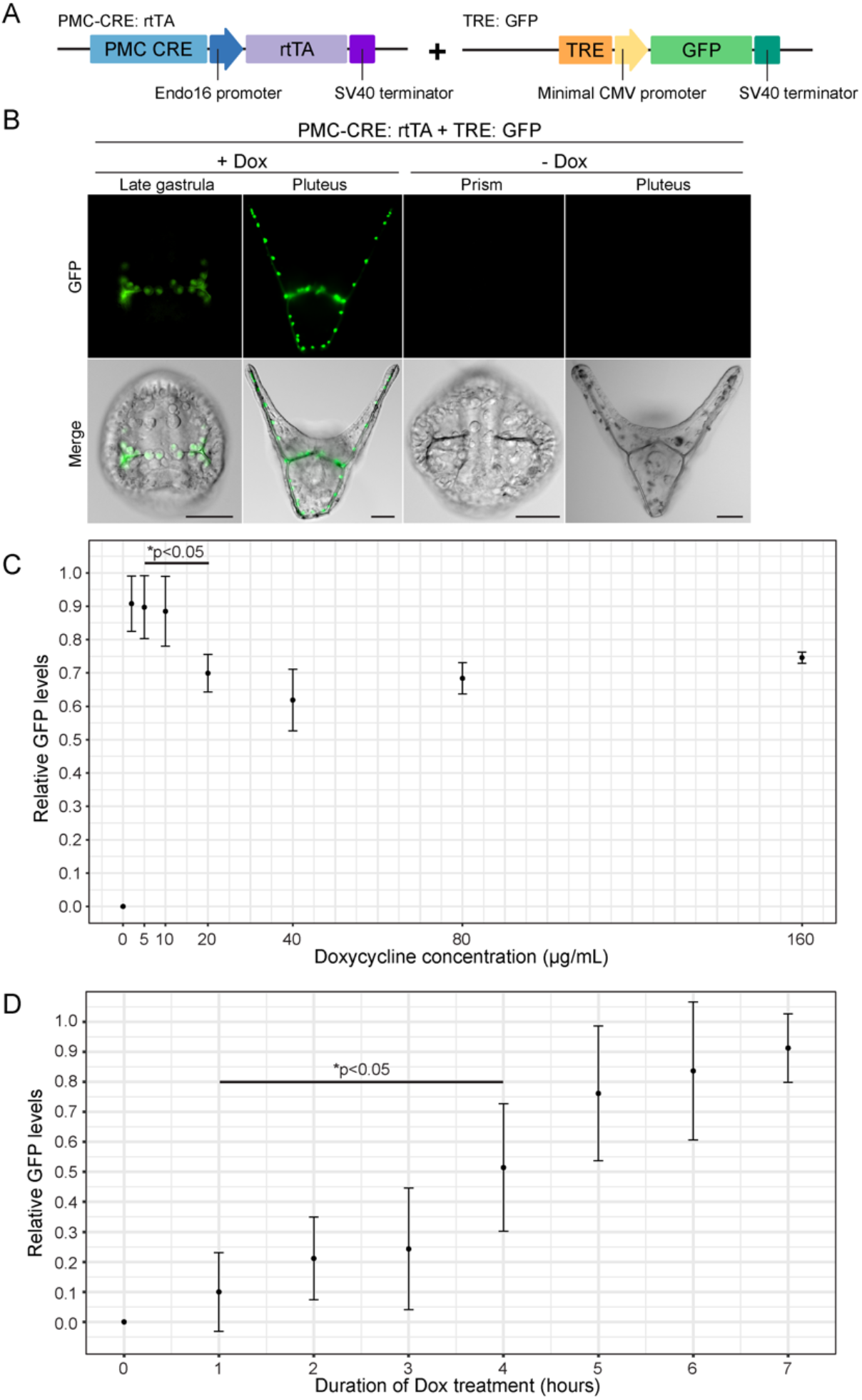
Inducible gene expression in the sea urchin embryo using the Tet-On system. (A) Schematic representation of the transactivator and responder constructs used to induce GFP expression in PMCs (see Materials and Methods). (B) GFP expression in the PMCs of transgenic embryos exposed to 5 μg/mL doxycycline (Dox). Top row: GFP fluorescence in live embryos. Bottom row: GFP fluorescence overlayed onto differential interference contrast (DIC) images. (C) A dose-response curve showing Dox dose-dependent induction of GFP expression. Embryos were treated with increasing concentrations of Dox (0, 2, 5, 10, 20, 40, 80, and 160 μg/mL) and relative GFP protein levels were quantified by ELISA (**see Figure S1A**). There was a moderate but statistically significant decrease in relative GFP levels (1.28-fold) when comparing embryos treated with 5 and 20 μg/mL Dox (see Supplemental Table S1). (D) Plot showing the time-dependent inducibility of GFP expression following Dox treatment. GFP expression was induced with 5 μg/mL Dox at the late gastrula stage and relative GFP protein levels were quantified by ELISA (**see Figure S1C,D**). There was a statistically significant increase in relative GFP levels (7.6-fold) at 4 hours compared to the first hour of Dox exposure (see Supplemental Table S1). Error bars show standard deviations from three independently repeated experiments. Scale bar: 50 μm.

### Optimization of doxycycline treatment

Although tetracycline and its derivatives such as doxycycline are generally well tolerated by eukaryotic systems, we asked whether Dox exposure can result in adverse effects during sea urchin embryonic development (**Figure S2A**). We found that embryos exposed to Dox at various concentrations and different stages developed normally (**Figure S2B**). We next asked whether Dox treatment can affect the development of transgenic embryos constitutively expressing rtTA (**Figure S3A**). Similarly, we found that transgenic embryos expressing PMC-specific rtTA.mCherry developed normally in the presence of Dox, regardless of the concentration or timing of exposure (**Figure S3B**). To examine the Dox dose-dependent response of our Tet-On system, we used ELISA to quantify GFP protein levels in transgenic embryos (PMC-CRE: rtTA + TRE: GFP) treated with various Dox concentrations (**Figure S1B**). Although the overall levels of responder inducibility varied between biological replicates (**Supplemental Table S1**), we found that induction of GFP expression was highly dose-dependent, with maximal activation achieved at Dox concentrations ranging from 2-10 μg/mL (**Figure 1C**). Based on these results, we chose to use 5 μg/mL of Dox for all our experiments, unless stated otherwise. We next evaluated the amount of time of Dox exposure that was needed for detectable gene expression in transgenic embryos co-injected with PMC-CRE: rtTA and TRE: GFP by measuring GFP protein levels in transgenic embryos exposed to 5 μg/mL Dox from 0-7 hours (**Figure S1C**). We found that a statistically significant increase in relative GFP levels (*p<0.05) could be detected via ELISA as early as 4 hours after the addition of Dox to the seawater (**Figure 1D**). We could also detect GFP expression by fluorescence imaging approximately 3-4 hours after the addition of Dox (**Figure S1D**). Taken together, our findings show that inducible GFP expression in sea urchin PMCs using the Tet-On system is dependent on the dose and duration of Dox exposure.

### PMC-specific induction of transgene expression in *L. variegatus* embryos

#### Dominant negative Ets1

To leverage the potential of our Tet-On system for studying sea urchin skeletogenesis, we sought to disrupt genes involved in the MAPK signaling pathway, an essential component of the GRN governing skeletogenic cell fate specification (**Figure S4**). Ets1, a downstream target of the MAPK signaling pathway, is a pivotal regulatory gene within the skeletogenic GRN and is required for PMC specification and epithelial-mesenchymal transition (EMT). The function of Ets1 during late skeletogenic processes, however, has not been explored. A dominant negative form of Ets1 consisting of only the DNA binding domain has been characterized in previous studies (Kurokawa et al., 1999; Sharma and Ettensohn, 2010). We modified this dominant form by fusing the coding sequences of the repressor domain of *Drosophila melanogaster* Engrailed (Dm-En) with the DNA-binding domain of *L. variegatus* Ets1 (Lv-Ets1-DBD), and tagged the chimeric protein with GFP, thereby creating dnLv-Ets1.GFP. We then cloned the recombinant gene downstream of the tetracycline response element (TRE) and minimal CMV promoter to generate the responder construct TRE: dnLv-Ets1.GFP (**Figure 2A**). To confirm the dominant negative effects of our chimeric protein, we synthesized and injected capped *dnLv-Ets1*.*GFP* mRNA into fertilized *L. variegatus* eggs. We confirmed that PMCs failed to ingress and skeletogenesis was inhibited in embryos expressing dominant negative Ets1, as reported in previous studies (**Figure S5A,B**). In contrast, embryos injected with mRNA encoding for a chimeric protein consisting of Dm-En and GFP (Dm-En.GFP) developed normally (**Figure S5C**).

**Figure 2:**
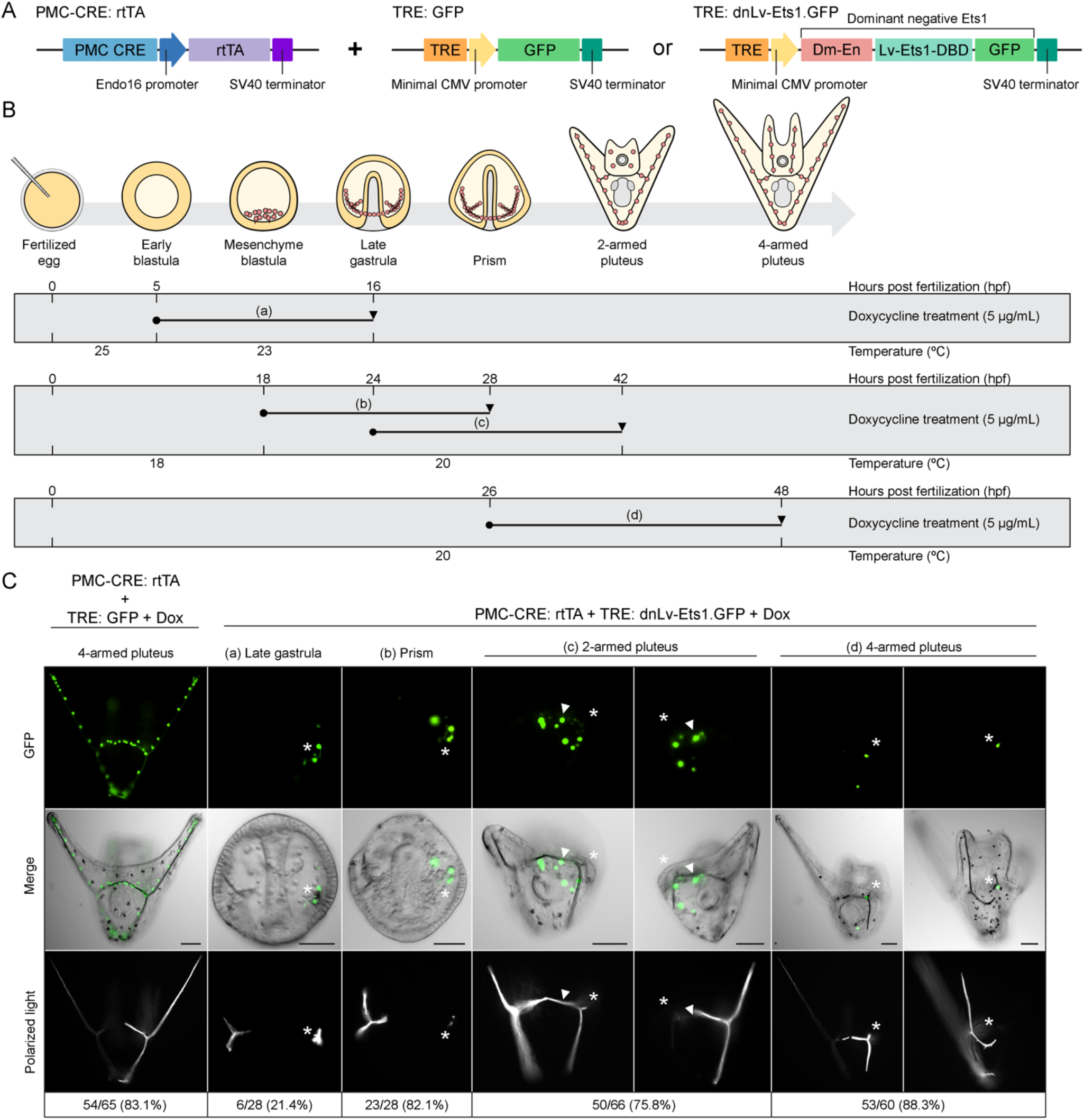
Localized expression of dominant negative Ets1 (dnLv-Ets1.GFP) in PMCs disrupts skeletogenesis in transgenic sea urchin embryos. (A) Schematic representation of the transactivator and responder constructs used to induce PMC-specific GFP or dnLv-Ets1-GFP expression. (B) Experimental design showing the treatment schedule and incubation temperatures (a-d). Solid circles indicate the stages at which doxycycline was added. Black arrowheads indicate the stages at which embryos were collected for analysis. (C) Representative images of transgenic embryos with induced expression of PMC-specific GFP or dnLv-Ets1-GFP (treatment schedule a-d). GFP expression in the PMCs did not affect embryonic skeletogenesis. Induced asymmetric expression of dnLv-Ets1-GFP in PMCs inhibited spicule formation and elongation of skeletal rods (asterisks). The proportion of transgenic embryos showing similar patterns of GFP or dnLv-Ets1-GFP expression and phenotype is shown. White arrowheads indicate ventral transverse rods that developed normally. Top row: GFP fluorescence in live embryos; Middle row: GFP fluorescence overlayed onto DIC images; Bottom row: Polarized light images showing the skeletal elements. Scale bar: 50 μm.

To induce dnLv-Ets1.GFP expression in PMCs, we co-injected the PMC-CRE: rtTA and TRE: dnLv-Ets1.GFP constructs into fertilized eggs. We exposed the transgenic embryos to Dox at different times and imaged them at various developmental stages (**Figure 2B**). We observed that dnLv-Ets1.GFP expression was restricted to specific PMC clonal clusters, likely due to the presence of a nuclear localization signal (NLS) within the Lv-Ets1 DNA-binding domain, coupled with mosaic incorporation of transgenes in sea urchin embryos during early cleavage stages. This finding was not surprising, as other transcription factors also exhibit highly restricted mobility within the PMC syncytium (Ettensohn et al., in preparation). Expression of dnLv-Ets1.GFP only on one side of the bilaterally symmetrical embryo allowed us to observe the distinct phenotype caused by perturbation of Ets1 function, as PMCs without transgene expression served as internal controls within the same individual embryos. When embryos were exposed to Dox from the early blastula stage and scored at the late gastrula stage, we found that only a small subset of transgenic embryos with asymmetric dnLv-Ets1.GFP expression exhibited unilateral defects in spicule formation (**Figure 2C**). This may be due to the low activity of the *Sp-EMI/TM* CRE during early development (i.e. prior to PMC ingression). Remarkably, however, at later stages of development, asymmetric expression of dnLv-Ets1.GFP almost always coincided with the inhibition of spicule formation or skeletal rod growth and elongation specifically on the side of the embryo where the dominant negative protein was expressed. Expression of dnLv-Ets1.GFP in PMCs did not appear to affect skeletal structures that had formed prior to induction of the transgene, such as the ventral transverse rods. Exposure to Dox prior to PMC ingression results in inhibition of spicule formation (**Figure 2C-a,b**). In contrast, embryos exposed to Dox during the late gastrula stage where PMCs have migrated exhibited shortened body and postoral rods (**Figure 2C-c**). When exposed to Dox at the prism stage, growth and elongation of the postoral and anterolateral rods were disrupted in 4-armed plutei (**Figure 2C-d**). These findings indicate that Ets1 is not only required for specifying PMCs early in development but also for actively maintaining proper growth and elongation of the skeletal rods.

To characterize the effects of dnLv-Ets1.GFP on the expression of terminal differentiation genes in the PMCs, we used ImmunoFISH to simultaneously visualize dnLv-Ets1.GFP protein and RNA transcripts of biomineralization genes (see Methods and Materials). Both *p16* and *sm30b* are downstream targets of Ets1, based on early knockdown of Ets1 expression and analysis of gene expression at the mesenchyme blastula stage (Rafiq et al., 2014). In control embryos, *p16* is highly expressed at the tips of the actively growing arm rods (the postoral and anterolateral rods) and at the tips of the body rods, in the scheitel region (**Figure 3A,B**). In contrast, the *sm30b* gene is highly expressed throughout the PMC syncytium, except for PMCs associated with the ventral transverse rods (**Figure 3A,C**). ImmunoFISH staining of transgenic embryos (PMC-CRE: rtTA + TRE: dnLv-Ets1.GFP) revealed that localized dnLv-Ets1.GFP expression in PMCs inhibited *p16* expression at the tip of the nearest arm (**Figure 3D**). Expression of dnLv-Ets1.GFP in the ventral transverse rods, however, did not appear to affect *p16* expression in the nearest arm of the 2-armed pluteus. Symmetric expression of dnLv-Ets1.GFP in the ventrolateral clusters of prism stage embryos also completely abolished *sm30b* expression (**Figure 3E**). Asymmetric, localized expression of dnLv-Ets1.GFP in transgenic embryos, however, abolished *sm30b* expression in PMCs only on that side of the bilateral 2-armed pluteus. The disruption to *sm30b* expression occurred not only in PMCs with detectable levels of dnLv-Ets1.GFP expression, but also PMCs nearby. These results show for the first time that positive inputs from Ets1 are required to maintain the expression of PMC terminal differentiation genes during the later stages of sea urchin embryonic development.

**Figure 3:**
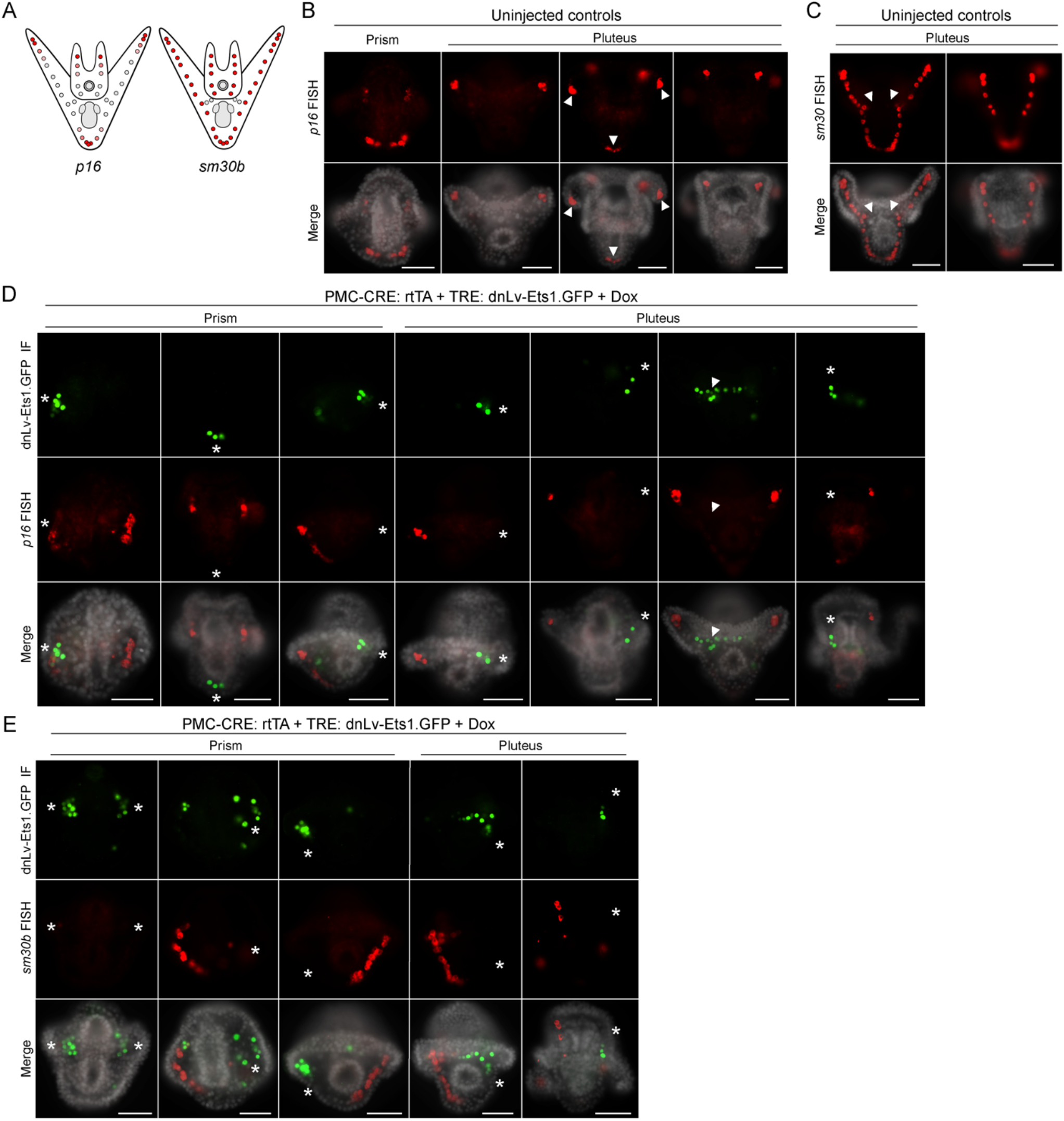
Localized expression of dominant negative Lv-Ets1 (dnLv-Ets1.GFP) in PMCs disrupts expression of downstream terminal differentiation genes, *p16* and *sm30b*. (A) Schematic representation of the expression patterns of *p16* and *sm30b* genes. (B) Single-color fluorescent in situ hybridization (FISH) images showing strong *p16* expression at the tips of the growing arms and the tips of the body rods (arrowheads). (C) Single-color FISH images showing strong *sm30b* expression throughout the PMC syncytial network but not in the ventral transverse rods (arrowheads). (D) GFP and *p16* immunoFISH staining of transgenic embryos with asymmetric dnLv-Ets1.GFP expression in PMCs. PMCs expressing dnLv-Ets1.GFP showed a reduction in *p16* expression in the nearest arm (asterisks). Expression of dnLv-Ets1.GFP in the ventral transverse rods did not affect *p16* expression in the growing arms of the 2-armed pluteus (arrowheads). Top row: GFP-immunostained cells; Middle row: Cy3-labeled *p16* RNA transcripts; Bottom row: Fluorescence merged with Hoechst 33342 counterstain in grayscale. (E) GFP and *sm30b* immunoFISH staining of transgenic embryos with asymmetric dnLv-Ets1-GFP expression in PMCs. PMCs expressing dnLv-Ets1.GFP show reduced *sm30b* expression. Expression of *sm30b* is also disrupted in PMCs nearby (asterisks). Top row: GFP-immunostained cells; Middle row: Cy3-labeled *sm30b* RNA transcripts; Bottom row: Fluorescence merged with Hoechst 33342 counterstain in grayscale.

Additionally, we also performed immunostaining with 6a9, a monoclonal antibody that reacts specifically with sea urchin MSP130 family of cell surface proteins, and anti-GFP antibody (**Figure S6A**). We found that while localized dnLv-Ets1.GFP expression disrupted the growth and elongation of skeletal elements (except for the ventral transverse rods), GFP-positive PMCs were still strongly labeled with 6a9. These findings suggest that protein degradation following disruption of the skeletogenic GRN may require more time to be observable, compared to the more rapid mRNA transcript turnover (**Figure S6B**).

#### MEK and DUSP6

Next, we sought to utilize the Tet-On system to induce expression of other genes within the MAPK signaling pathway, which has been shown to regulate Ets1 activity at early developmental stages (Röttinger et al., 2004). We chose to focus on the mitogen-activated protein kinase kinase (MEK) and dual specificity phosphatase 6 (DUSP6), which are enzymes that regulate the extracellular signal-regulated kinase (ERK) (Guo et al., 2020; Muhammad et al., 2018). MEK directly phosphorylates ERK to activate the MAPK signaling pathway while DUSP6 directly dephosphorylates ERK to negatively regulate signaling. The amino acid sequences of the sea urchin MEK and DUSP6 proteins share a high degree of conservation with their human orthologs (**Figure S7**). Unlike dnLv-Ets1.GFP, MEK and DUSP6 do not strictly localize to the nucleus (**Figure 4 and Figure 5**); suggesting that these proteins might translocate and exert their functions throughout the PMC syncytial network.

**Figure 4:**
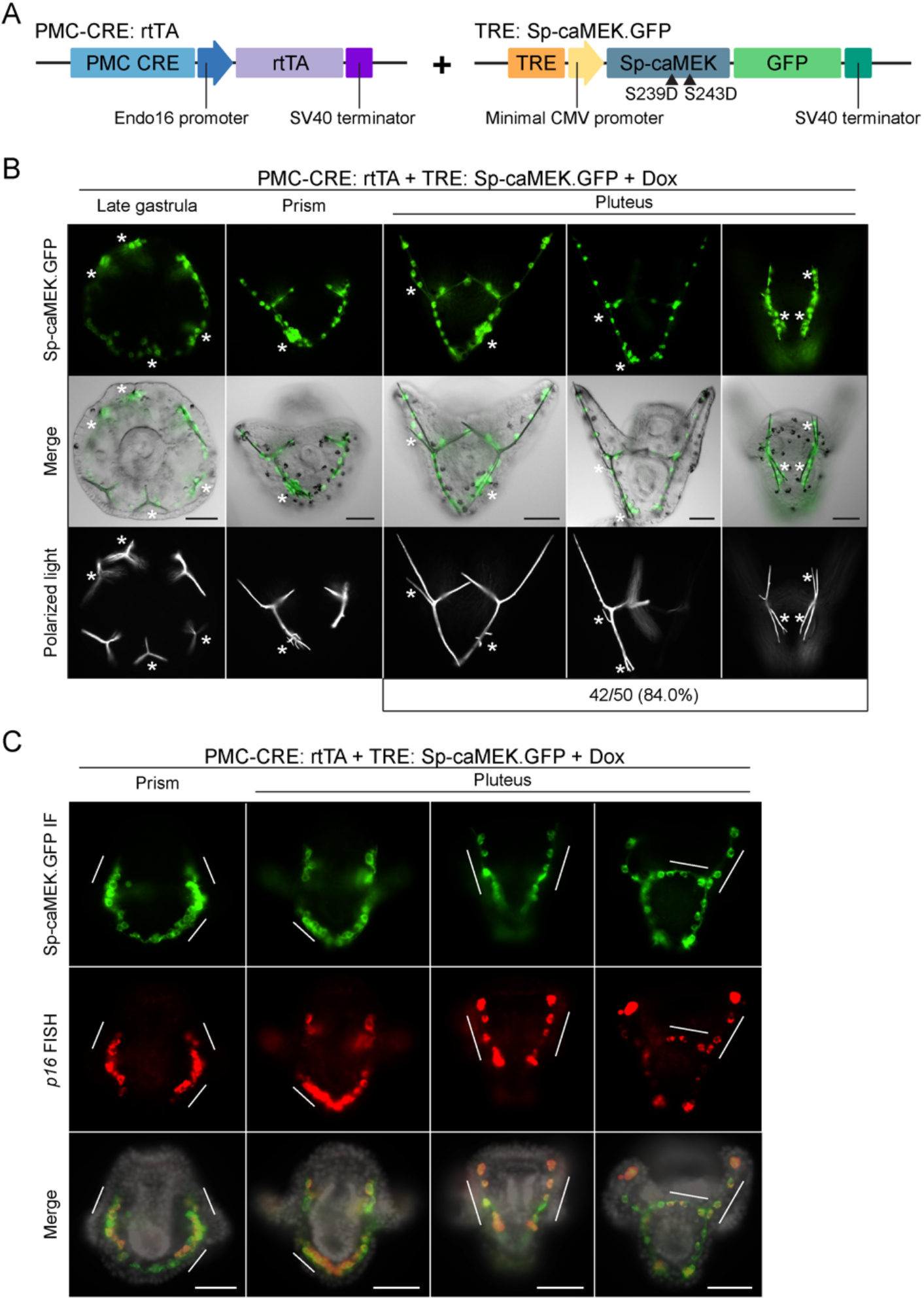
Induced expression of constitutively active *S. purpuratus* MEK (Sp-caMEK.GFP) in PMCs disrupts skeletal patterning. (A) Schematic representation of the transactivator and responder constructs used to induce Sp-caMEK.GFP expression in PMCs. (B) Images of transgenic embryos expressing Sp-caMEK.GFP. The protein is distributed throughout the PMC syncytial network, resulting in supernumerary spicules and abnormal skeletal branching (asterisks). The number of embryos with Sp-caMEK.GFP expression that showed the abnormal skeletal branching phenotype is indicated. Top row: GFP fluorescence in live embryos; Middle row: GFP fluorescence overlayed onto DIC images; Bottom row: Polarized light images showing skeletal elements. (C) GFP and *p16* immunoFISH staining of transgenic embryos expressing Sp-caMEK.GFP showed expansion of the *p16* expression domain (white bars). Top row: GFP-immunostained cells; Middle row: cy3-labeled *p16* RNA transcripts; Bottom row: Fluorescence merged with Hoechst 33342 counterstain in grayscale. Scale bar: 50 μm.

**Figure 5:**
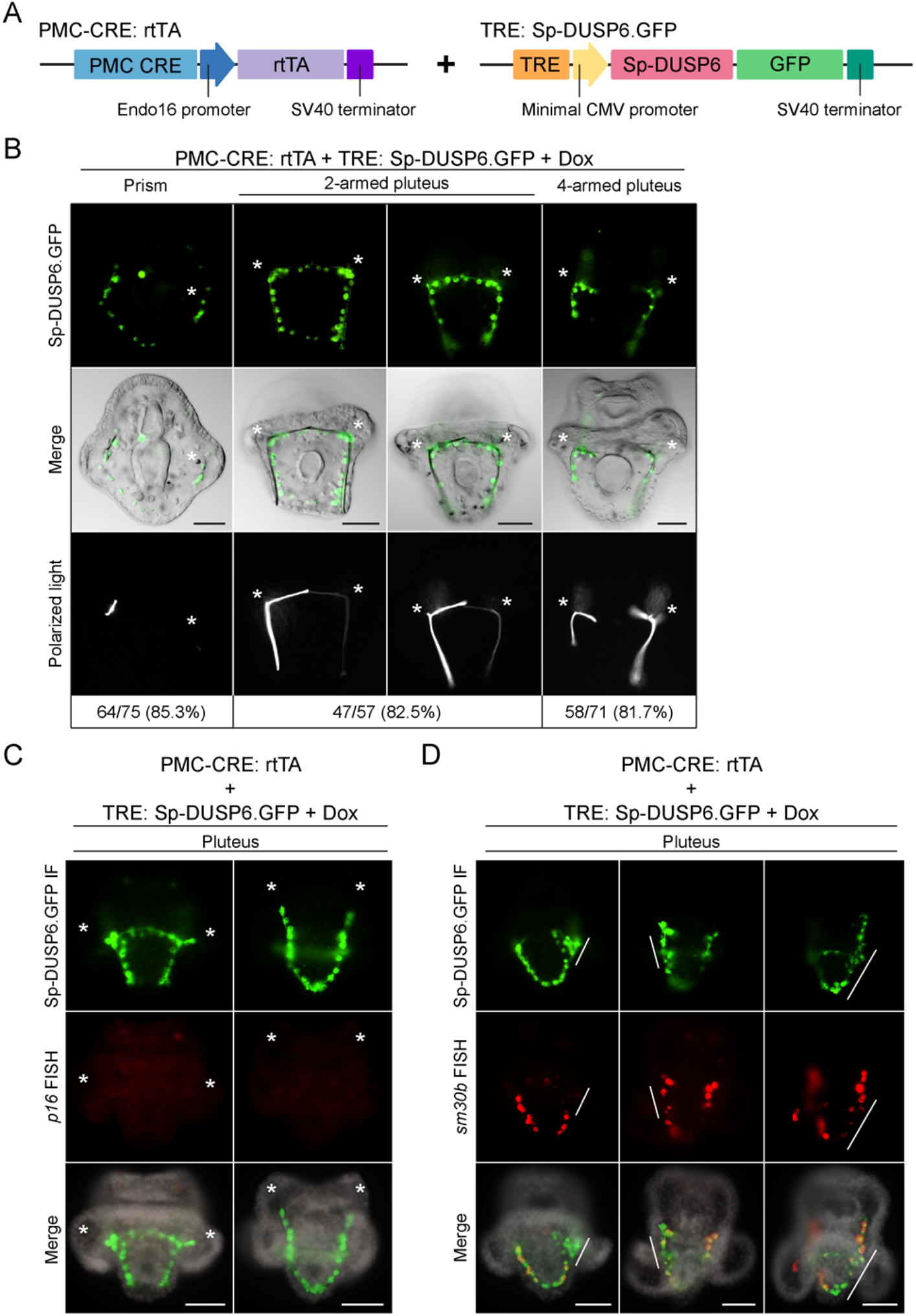
Induced expression of *S. purpuratus* DUSP6 (Sp-DUSP6.GFP) in PMCs inhibits skeletogenesis. (A) Schematic representation of the transactivator and responder constructs used to induce Sp-DUSP6.GFP expression in PMCs. (B) Representative images of transgenic embryos expressing Sp-DUSP6.GFP. The protein is distributed throughout the PMC syncytial network, inhibiting spicule formation and skeletal growth (asterisks). The numbers of embryos with Sp-DUSP.GFP expression that showed an abnormal skeletal growth and elongation phenotype were scored. Top row: GFP fluorescence in live embryos; Middle row: GFP fluorescence overlayed onto DIC images; Bottom row: Polarized light images showing skeletal elements. (C) GFP and *p16* immunoFISH staining of transgenic embryos expressing Sp-DUSP6-GFP in PMCs showing loss of *p16* expression (asterisks). Top row: GFP-immunostained cells; Middle row: Cy3-labeled *p16* RNA transcripts; Bottom row: Fluorescence merged with Hoechst 33342 counterstain in grayscale. (D) GFP and *sm30b* immunoFISH staining of transgenic embryos expressing Sp-DUSP6.GFP showing partial loss of *sm30b* expression in some PMCs (white bars). Top row: GFP-immunostained cells; Middle row: Cy3-labeled *sm30b* RNA transcripts; Bottom row: Fluorescence merged with Hoechst 33342 counterstain in grayscale. Scale bar: 50 μm.

To investigate the effects of MEK overexpression in sea urchin embryos, we cloned a constitutively active form of *S. purpuratus* MEK (S239D/S243D) into the responder construct to generate TRE: Sp-caMEK.GFP (**Figure 4A**). The phosphomimetic mutations that we introduced in Sp-caMEK are located in a highly conserved domain that is also found in human MEK1 (**Figure S7A**). We first injected capped mRNA containing the coding sequence of Sp-caMEK.GFP into fertilized *L. variegatus* eggs. We found that overexpression of Sp-caMEK.GFP resulted in the formation of ectopic skeletal spicules as well as abnormal branching of skeletal rods (**Figure S8A,B**). To induce Sp-caMEK.GFP expression, we co-injected the PMC-CRE: rtTA and TRE: Sp-caMEK.GFP constructs into fertilized eggs. Upon overnight exposure to Dox, the Sp-caMEK.GFP protein was found to be present in all PMCs. We found that late gastrula stage embryos with Sp-caMEK.GFP expression exhibited ectopic spicule formation (**Figure 4B**). In prism and pluteus embryos that were treated with Dox overnight, Sp-caMEK.GFP expression resulted in abnormal branching of the skeletal rods. Using immunoFISH, we also found that Sp-caMEK.GFP overexpression expanded the spatial expression domain of the *p16* gene beyond PMCs that were located near the tips of the growing arms and body rods (**Figure 4C**).

We next injected capped mRNA containing the coding sequence of Sp-DUSP6.GFP into fertilized *L. variegatus* eggs. As previously reported by Röttinger et al., 2004, we found that injection of Sp-DUSP6.GFP mRNA into fertilized eggs completely inhibited PMC specification and spicule formation (**Figure S8C,D**). We then cloned Sp-DUSP6 into the responder construct to generate TRE: Sp-DUSP6.GFP (**Figure 5A**). In prism stage transgenic embryos that were treated with Dox overnight, Sp-DUSP6.GFP overexpression inhibited spiculogenesis (**Figure 5B**). In pluteus stage embryos that were treated with Dox overnight, Sp-DUSP6.GFP overexpression completely abolished the growth and elongation of the skeletal rods. The Sp-DUSP6.GFP protein was also observed to have translocated throughout the PMC syncytium. ImmunoFISH of transgenic embryos with induced Sp-DUSP6.GFP expression revealed that *p16* expression was completely abolished (**Figure 5C**), while *sm30b* expression was partially lost in a subset of PMCs (**Figure 5D**). Taken together, our findings provide additional evidence that MAPK signaling is essential for the non-uniform patterns of expression within the PMC syncytium following the shift in cell-autonomous to signal-dependent regulation of the skeletogenic GRN. Our observations are consistent with previous studies showing that exposure to a MEK inhibitor (U0126) at later stages inhibited skeletogenesis (Fernandez-Serra et al., 2004; Röttinger et al., 2004). Additionally, these data further support the usefulness of the Tet system for probing the functions of pathways at late developmental stages in situations where inhibitors are not available, with the added advantage of cellular specificity.

### Cell type-specific induction of transgene expression in *S. purpuratus* embryos

As many validated CREs have been discovered and tested in *S. purpuratus*, we asked whether we could use our two-plasmid Tet-On system to drive GFP expression in this species. In the same studies, we also sought to confirm that the Tet-On system could be used to drive gene expression in cell types other than PMCs. In initial experiments, we co-injected the PMC-CRE: rtTA and TRE: GFP constructs into fertilized *S. purpuratus* eggs. Transgenic embryos exposed to Dox at the late gastrula stage showed no observable GFP expression at the early pluteus stage (data not shown). As the PMC CRE we used was originally derived from *S. purpuratus* and has been shown to drive robust, PMC-specific gene expression in both *S. purpuratus* and *L. variegatus*, this finding pointed to a potential species-specific limitation of the particular pair of transactivator and responder plasmids. To overcome this hurdle, we tested a different combination of transactivator and responder constructs (see Materials and Methods) (**Figure 6A**). Surprisingly, transgenic embryos co-injected with the PMC CRE: tetON3G and TRE3Gp: GFP constructs were able to show strong PMC-specific GFP expression upon overnight exposure to Dox (**Figure 6B**). We also co-injected the same pair of constructs into *L. variegatus* embryos and found that they were similarly functional (**Figure 6C**).

**Figure 6:**
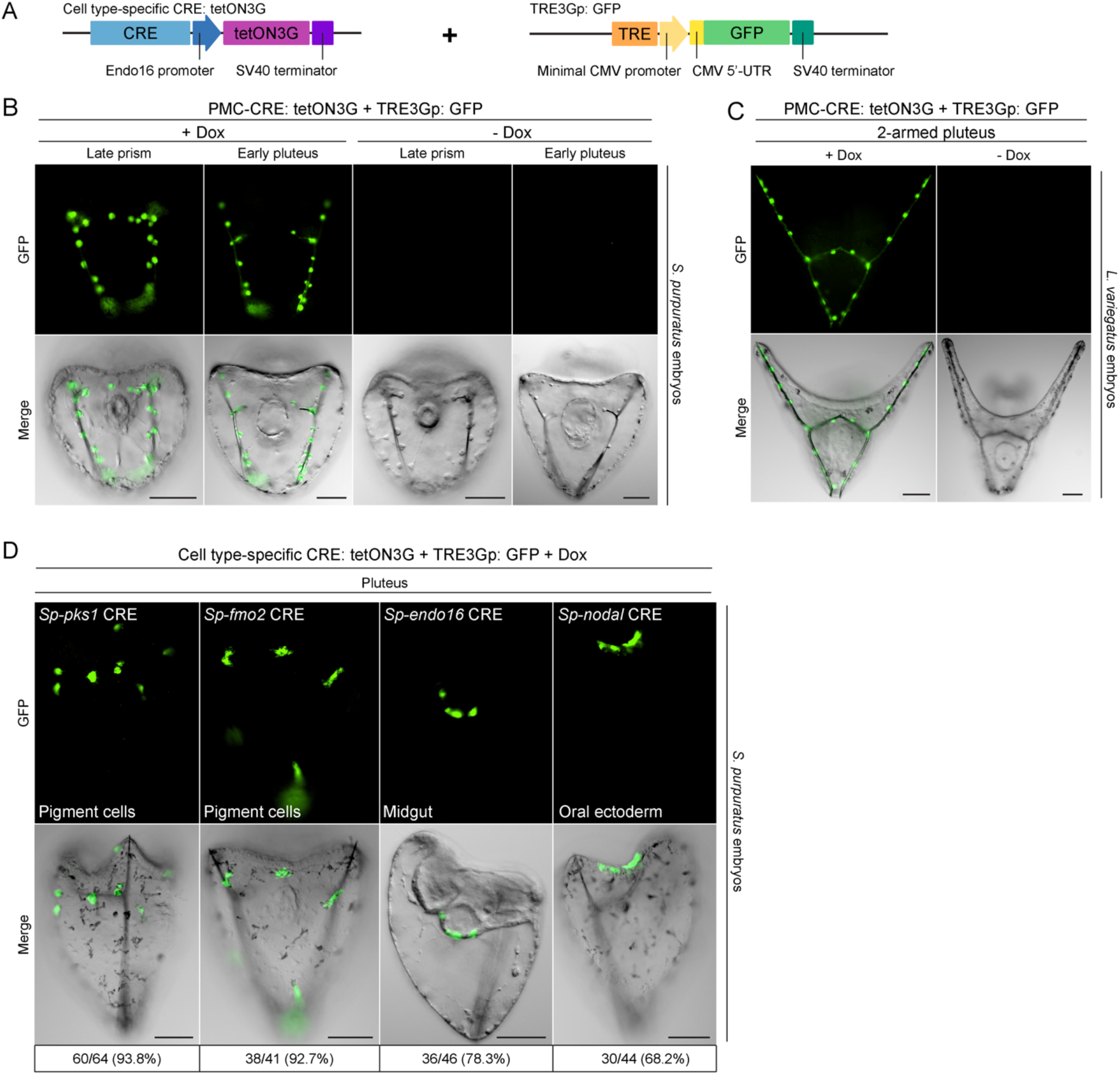
Inducible gene expression in diverse cell types using the Tet-On system. (A) Schematic representation of the transgenic activator and responder constructs used to induce GFP expression in *S. purpuratus* embryos (see Materials and Methods). (B, C) GFP expression in the PMCs of transgenic *S. purpuratus* and *L. variegatus* embryos exposed to 5 μg/mL Dox at the late gastrula stage. GFP fluorescence was not observed when embryos were not exposed to Dox. Top row: GFP fluorescence in live embryos. Bottom row: GFP fluorescence overlayed onto differential interference contrast (DIC) images. (D) GFP expression in different cell types of transgenic embryos exposed to 5 μg/mL Dox at the late gastrula stage. The number of embryos with GFP fluorescence showing expression in the expected cell types was scored. Top row: GFP fluorescence in live embryos; Bottom row: GFP fluorescence overlayed onto differential interference contrast (DIC) images. Scale bar: 50 μm.

To take further advantage of the plethora of validated *S. purpuratus* CREs available, we generated tetON3G transactivator constructs containing CREs that can drive cell type-specific expression in the pigment cells (*Sp-pks1* and *Sp-fmo2*) (Khor et al., 2021), gut (*Sp-endo16*) (Yuh and Davidson, 1996), and oral ectoderm (*Sp-nodal*) (Nam et al., 2007). We found that we were able to induce cell type-specific GFP expression upon overnight exposure to Dox (**Figure 6D**). Taken together, these results emphasize the wide utility of our Tet-On system as a tool for inducible, cell type-specific gene expression in sea urchins.

### Tet-On system for nitroreductase-mediated cell ablation

Targeted cell ablation is a powerful tool for investigating the *in vivo* function of cells. Previous studies have analyzed cell-cell interactions in living sea urchin embryos using fluorescence photoablation (Ettensohn, 1990). Here, we developed a dual input Tet-On system for nitroreductase-mediated cell ablation in sea urchin embryos. Bacterial nitroreductase sensitizes cells to metronidazole (MTZ) by converting the prodrug into a cytotoxic product (Lindmark and Müller, 1976). We cloned a rationally engineered nitroreductase ortholog (NTR 2.0) from *Vibrio vulnificus* (Sharrock et al., 2022) into the Tet-On responder construct to generate TRE: NTR-2.0.GFP (**Figure 7A**). We then co-injected the PMC-CRE: rtTA and TRE: NTR-2.0.GFP constructs into fertilized *L. variegatus* eggs. Transgenic embryos were first exposed to Dox at the early blastula stage (**Figure 7B**). At the late gastrula stage, a subset of the transgenic embryos was treated with MTZ overnight. We found that overexpression of NTR 2.0 without MTZ exposure did not affect embryonic development (**Figure 7C**). In contrast, exposing transgenic embryos with NTR-2.0.GFP expression to MTZ resulted in PMC cell death as well as disruption to the growth and elongation of the pluteus arms and skeletal rods. We also observed that MTZ treatment did not affect transgenic sea urchin embryo development without NTR-2.0.GFP expression. Immunostaining of a PMC cell surface marker (MSP130) and GFP revealed that the expression of NTR-2.0.GFP, in combination with MTZ treatment, resulted in PMC ablation and disordered PMC syncytial cables (**Figure 7D**). The dual-input requirement of this system allows for improved control of targeted cell ablation in two steps: (1) Dox induction permits NTR-2.0 GFP to accumulate within cells, and (2) subsequent addition of MTZ causes rapid ablation as cells are already primed with the enzyme.

**Figure 7:**
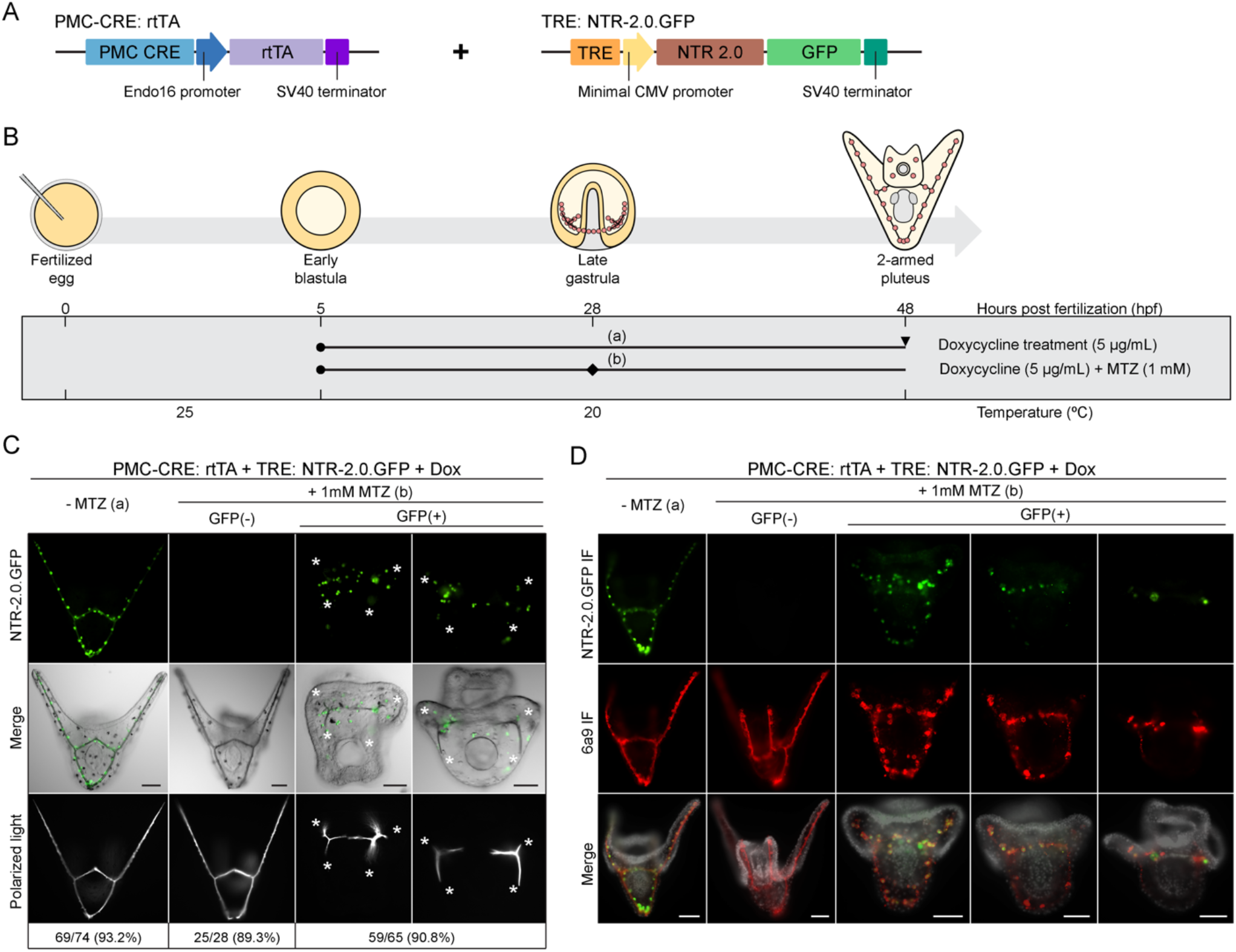
Dual input Tet-On system for nitroreductase-mediated cell ablation in sea urchin embryos. (A) Schematic representation of the transactivator and responder constructs used to induce nitroreductase (NTR-2.0.GFP) expression in PMCs. (B) Experimental design showing the treatment schedule and incubation temperatures. Solid circles indicate the stage at which doxycycline was added. Diamond represents the stage at which metronidazole (MTZ) was added. Arrowhead indicates the stage at which embryos were collected for analysis. (C) Induced expression of NTR-2.0.GFP and the addition of MTZ resulted in targeted ablation of PMCs and inhibition of skeletogenesis (asterisks). Top row: GFP fluorescence in live embryos; Middle row: GFP fluorescence overlayed onto DIC images; Bottom row: Polarized light images showing skeletal elements. (D) PMC marker (6a9) and GFP immunostaining of transgenic embryos expressing NTR-2.0.GFP in PMCs. Expression of NTR-2.0.GFP and in combination with MTZ treatment disrupted the PMC syncytial cables. Top row: GFP-immunostained cells; Middle row: 6a9-immunostained skeletal structures; Bottom row: Fluorescence merged with Hoechst 33342 counterstain in grayscale. Scale bar: 50 μm.

## Discussion

Sea urchins and other echinoderms are powerful models for studying developmental processes and the GRNs that govern them. There are diverse tools available for manipulating gene expression in echinoderm embryos. For instance, most functional studies involve the microinjection of morpholino antisense oligonucleotides (MOs), which often leads to specific and efficient gene knockdowns. Another widely used approach is the microinjection of mRNAs, typically encoding dominant negative or constitutively active proteins. As MOs and mRNAs are usually microinjected into fertilized eggs, a major limitation of these perturbations is that they are uncontrolled spatially and temporally; i.e., they affect all cells of the embryo from the onset of development until the reagent (MO or mRNA) declines in abundance to non-functional levels. Chemical inhibitors can partly overcome this hurdle by allowing control over the timing of their application. Not all molecules of pathways can be targeted by inhibitors, however, and inhibitors can produce unintended side effects, due to a lack of specificity or the pleiotropic nature of their targets.

As many sea urchin genes show dynamic changes in their spatial-temporal expression patterns during development, the inability to conditionally perturb them in a cell type-specific manner is a major obstacle in investigating the late developmental functions of genes, especially those that have crucial roles during early embryogenesis. With respect to GRN biology, the lack of targeted approaches has made it difficult to analyze dynamic changes in network circuitry; for example, to determine whether a regulatory gene with an important early function continues to provide regulatory inputs at later developmental stages or, alternatively, hands off its regulatory function to downstream transcription factors. In addition, as most regulatory genes are expressed in multiple territories, embryo-wide perturbations can lead to significant inaccuracies in the construction of cell type-specific GRN models. The ability to regulate gene perturbations in sea urchins would make it possible for the direct interrogation of GRN circuitry at late developmental stages and/or in specific tissues.

In the present study, we developed and optimized a two-plasmid Tet-On system for inducible gene expression in sea urchin embryos. In line with our goal to induce transient expression of our genes of interest, we opted to use the Tet-On system rather than Tet-Off, as the latter would require long-term exposure to doxycycline (Dox) to prevent constitutive expression. As proof of concept, we demonstrated the feasibility of the system by inducing the expression of GFP exclusively in the PMCs. It should be noted that the level of induction is expected to be dependent on different factors such as the activity of the CRE controlling rtTA expression, folding and maturation kinetics of the proteins, and half-life of the mRNA and protein products.

We next sought to expand the utility of the Tet-On system by using it to induce the expression of genes involved in the MAPK signaling pathway, a pivotal regulator of the sea urchin skeletogenic GRN. Two major components of the pathway, ERK and Ets1, are expressed in several cellular territories across a broad range of developmental stages (Fernandez-Serra et al., 2004; Röttinger et al., 2004). Hence, they are of special interest as candidates for possessing late developmental roles. Ets1 is a key transcription factor required for early PMC specification, and the Ets1 DNA-binding domain (DBD) has been shown to exert dominant negative functions when overexpressed (Kurokawa et al., 1999; Sharma and Ettensohn, 2010). Although the mechanism behind this effect is not entirely clear, we postulate that overexpression of the Ets1 DBD that lacks the transactivation domain outcompetes endogenous Ets1 for DNA binding sites. In this study, we fused the *L. variegatus* Ets1 DBD to the repressor domain of *Drosophila* Engrailed (Margolin et al., 1994), which efficiently converted the chimeric protein into an obligate repressor (dnLv-Ets1.GFP). We determined the pivotal role of Ets1 in regulating late stage skeletogenesis by inducing dnLv-Ets1.GFP expression at different developmental stages. Mosaic transgenesis in the present study offered a unique advantage when expressing a nuclear protein such as dnLv-Ets1.GFP, which exhibits limited mobility within the PMC syncytium. For instance, it was possible to observe the distinct phenotype caused by disruption of Ets1 function on one side of the bilaterally symmetrical embryo, while PMCs without transgene expression served as internal controls within the same embryos. We also observed that dnLv-Ets1.GFP locally disrupted the expression of biomineralization genes only in the vicinity of the transgenic cells. Significantly, dnLv-Ets1.GFP expression appeared to only inhibit skeletal elements that were actively growing at the time of induction. We observed that the late function of Ets1 was restricted to the postoral and anterolateral rods in 2-armed and 4-armed plutei, where VEGF3 is highly expressed by the adjacent ectoderm (Adomako-Ankomah and Ettensohn, 2013; Duloquin et al., 2007). In mammalian systems, VEGF stimulates the MAPK signaling pathway, thereby regulating cell proliferation and differentiation (Doanes et al., 1999). While both inputs are required for sea urchin skeletogenesis, a direct association between VEGF and MAPK signaling has not been established. Our analysis has for the first time directly probed the regulatory circuitry of the PMC GRN at post-blastula stages, when signals from VEGF and the MAPK pathway locally regulate skeletal growth and has shown that Ets1 provides essential, late inputs into biomineralization genes. These findings are consistent with the hypothesis that the VEGF and MAPK pathways act through Ets1, which at early developmental stages is regulated positively by MAPK signaling.

Activated ERK is expressed by many different groups of cells during sea urchin development. Previous studies have reported that MAPK signaling is required for PMC and pigment cell specification, skeletogenesis, and gut formation (Fernandez-Serra et al., 2004; Röttinger et al., 2004). Due to the diverse tissue and developmental processes regulated by MAPK, the common approach of using chemical inhibitors makes it challenging to pinpoint the direct role of MAPK signaling in regulating sea urchin skeletogenesis. Significantly, it has not been possible to determine whether ERK/MAPK signaling acts cell-autonomously within PMCs to regulate skeletogenesis or indirectly through effects on other tissues. Using the Tet-On system to induce expression of constitutively active MEK (Sp-caMEK.GFP) or DUSP6 (Sp-DUSP6.GFP) exclusively in PMCs, we were able to target ERK activity in a cell type-specific manner. We observed that embryos overexpressing Sp-caMEK.GFP exhibited dramatic, ectopic skeletal branching. While ectopic Sp-caMEK.GFP expression resulted in upregulation of *p16* throughout the PMC syncytium, we observed slightly elevated expression of *p16* at the tips of the growing arms, where it is normally expressed (Sun and Ettensohn, 2014). In contrast, ectopic Sp-DUSP6.GFP overexpression phenocopied late stage U0126 treatment and we observed only a partial loss of *sm30b* expression. These findings raise the possibility that factors other than ERK activity are involved in regulating downstream biomineralization genes such as *p16* and *sm30b*, although an alternative interpretation is that these molecular perturbations only partially disrupted ERK signaling.

It is widely thought that the embryonic skeleton arose within echinoderms via co-option of the adult skeletogenic program, as evidenced by the many similarities in the GRN of skeletogenic cells in the embryo and adult (Czarkwiani et al., 2013; Gao and Davidson, 2008; Gao et al., 2015; Killian et al., 2010). Significantly, the work we describe here allows for higher resolution dissection of the embryonic skeletogenic GRN by decoupling the early, cell-autonomous, and late, signal-dependent modes of skeletogenesis. As the MAPK signaling pathway is activated cell-autonomously in the early embryo, possibly through the actions of localized maternal factors such as β-catenin (Fernandez-Serra et al., 2004; Röttinger et al., 2004), we proposed that the heterochronic shift in the deployment of the skeletogenic GRN that occurred during euechinoid evolution was achieved by an important evolutionary innovation that placed MAPK signaling under maternal control. We favor a model whereby current regulatory mechanisms that underlie late embryonic skeletal patterning and growth reflect an ancient heterochronic shift in VEGF3 expression by ectodermal cells (Morino et al., 2012; Yamazaki et al., 2021). We hypothesize that VEGF3 activates Ets1 through the MAPK signaling pathway at late embryonic stages, circuitry that is likely to be similar to the ancestral skeletogenic GRN found in the adult.

To confirm the broader utility of the Tet-On system in sea urchins, we applied it to a second species, *S. purpuratus*. In preliminary experiments, we found that the transactivator and responder constructs used in *L. variegatus* embryos were not functional in *S. purpuratus*. We fully resolved this issue by using a different combination of constructs (TetON3G and TRE3Gp). The rtTA-advanced and TetON3G transactivators are distinct by only three amino acid changes while the TRE3Gp responder contains an additional CMV 5’-UTR downstream of the promoter. We showed that the new combination of transactivator and responder is also highly effective in *L. variegatus* embryos. Thus, in the absence of additional information regarding possible species-specificity of the expression system, we recommend that the TetON3G and TRE3Gp constructs be used in future applications. Using validated *S. purpuratus* CREs, we further showed that the Tet-On system can be used to induce gene expression in diverse cell types, a demonstration of the capabilities of the system. In the future, the Tet-On system will serve as a flexible platform for the spatio-temporal regulation of gene expression in sea urchins and other echinoderms, thereby enhancing the utility of these model organisms for developmental studies.

## Acknowledgments

We are grateful to Josiah Saunders for synthesizing the RNA probes used in this study. This work was supported by grants from the National Institutes of Health (R24-OD023046) and the National Science Foundation (IOS2004952), both to C.A.E.

---

**Figure S1:**
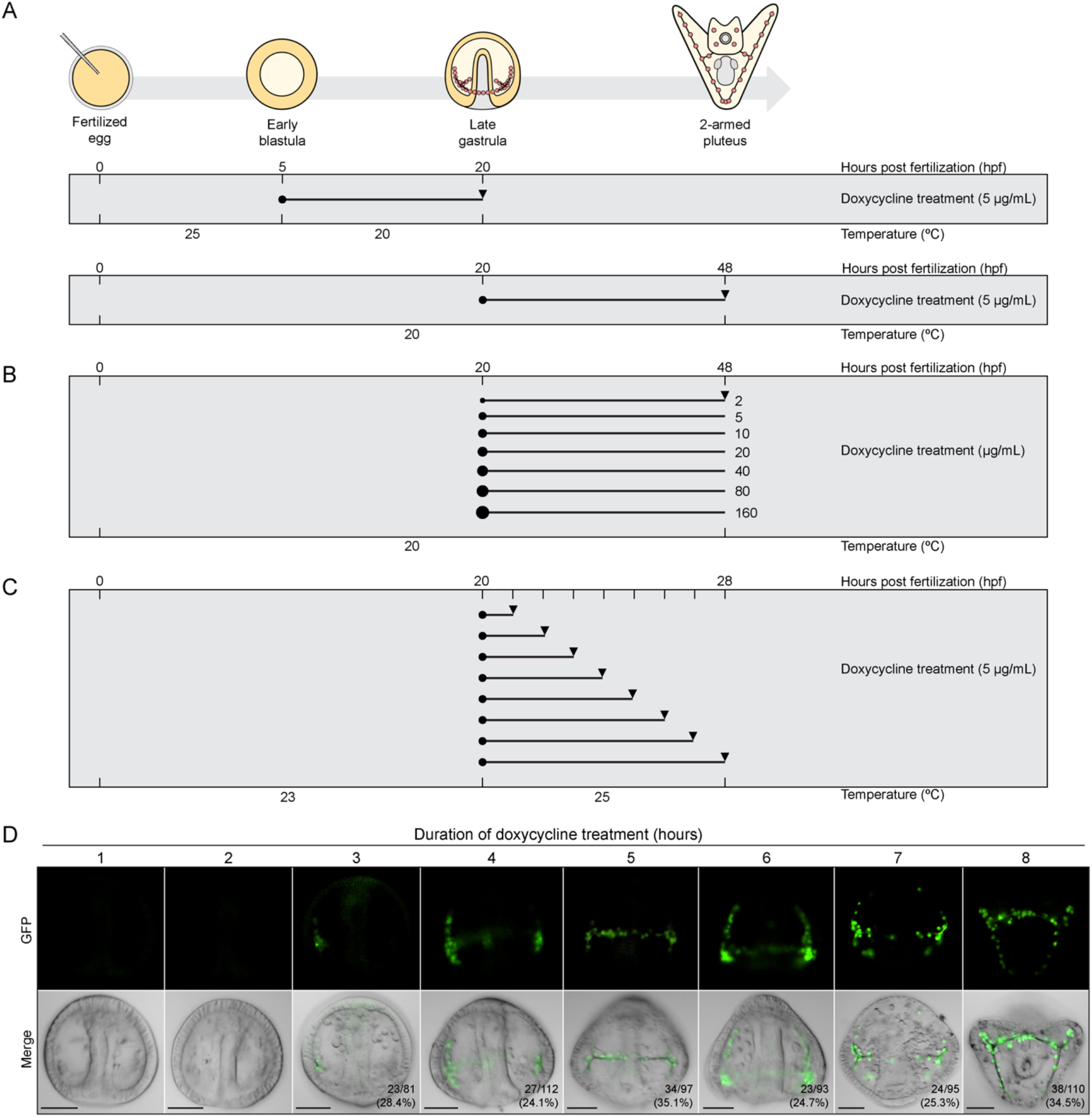
Doxycycline dose- and time-dependent induction of GFP expression. (A-C) Schematic representation of the experimental design showing the treatment schedule and embryo incubation temperatures. Experimental designs for Figure 1C (B) and Figure 1D (C) are shown. Solid circles indicate the stages at which Dox was added. Arrowheads indicate the stages at which embryos were collected for analysis. (D) Time-dependent induction of PMC-specific GFP expression in transgenic embryos treated with 5 μg/mL Dox. Dox treatment began at the late gastrula stage and embryos were then cultured continuously in the drug until they were collected for analysis. Injected embryos that showed GFP expression in PMCs were scored. Top row: GFP fluorescence in live embryos; Bottom row: GFP fluorescence overlayed onto DIC images. Scale bar: 50 μm.

**Figure S2:**
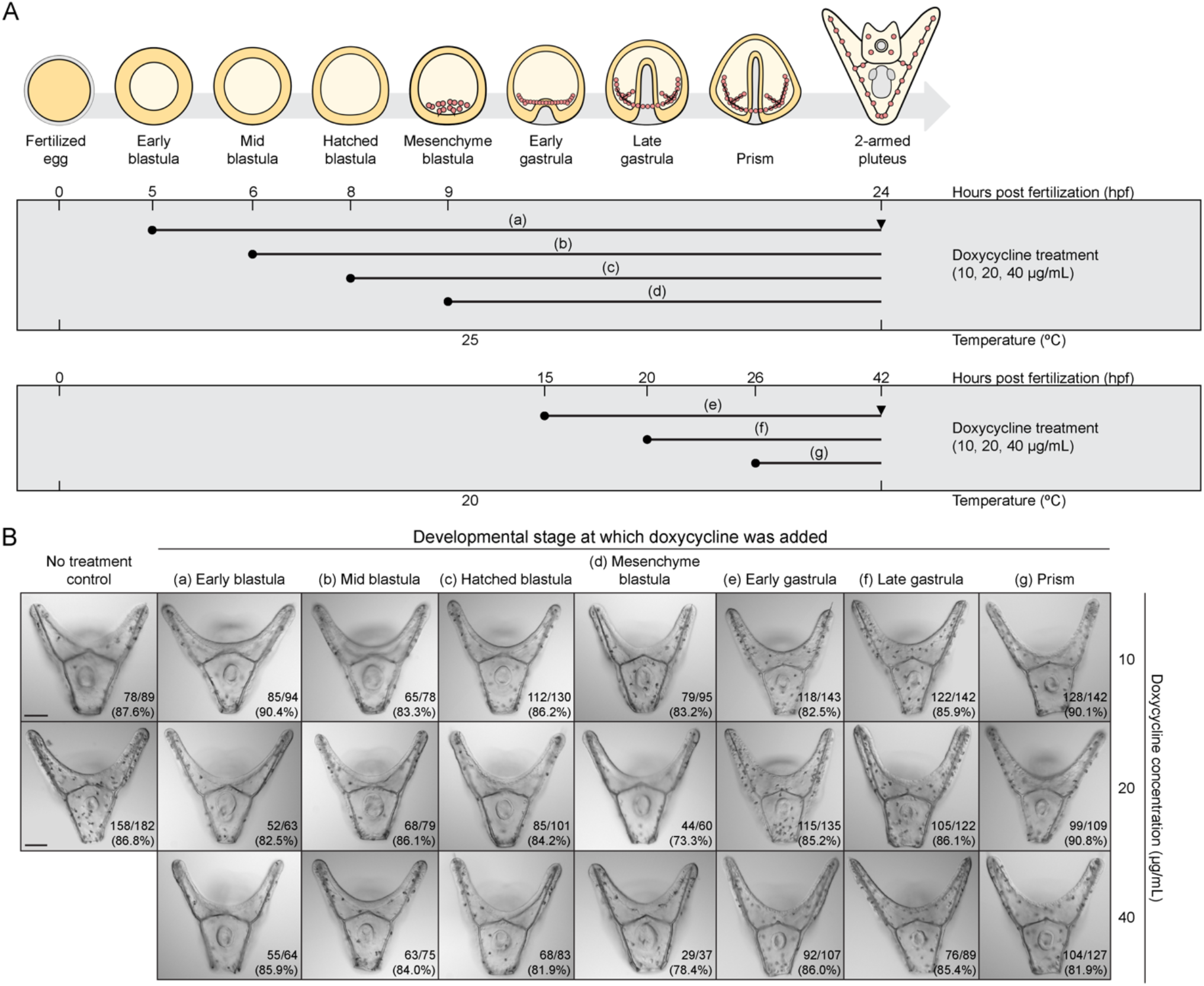
Development is not affected by doxycycline. (A) Schematic representation of the experimental design showing the treatment schedule and incubation temperatures. Dox at concentrations of 10, 20, and 40 μg/mL was added to uninjected control embryo cultures at various developmental stages (a-g). Solid circles indicate the stages at which doxycycline was added. Arrowheads indicate the stages at which embryos were collected for analysis. (B) Representative DIC images of live 2-armed plutei treated with Dox. The number of healthy embryos that reached the 2-armed pluteus stage was scored. Scale bar: 50 μm.

**Figure S3:**
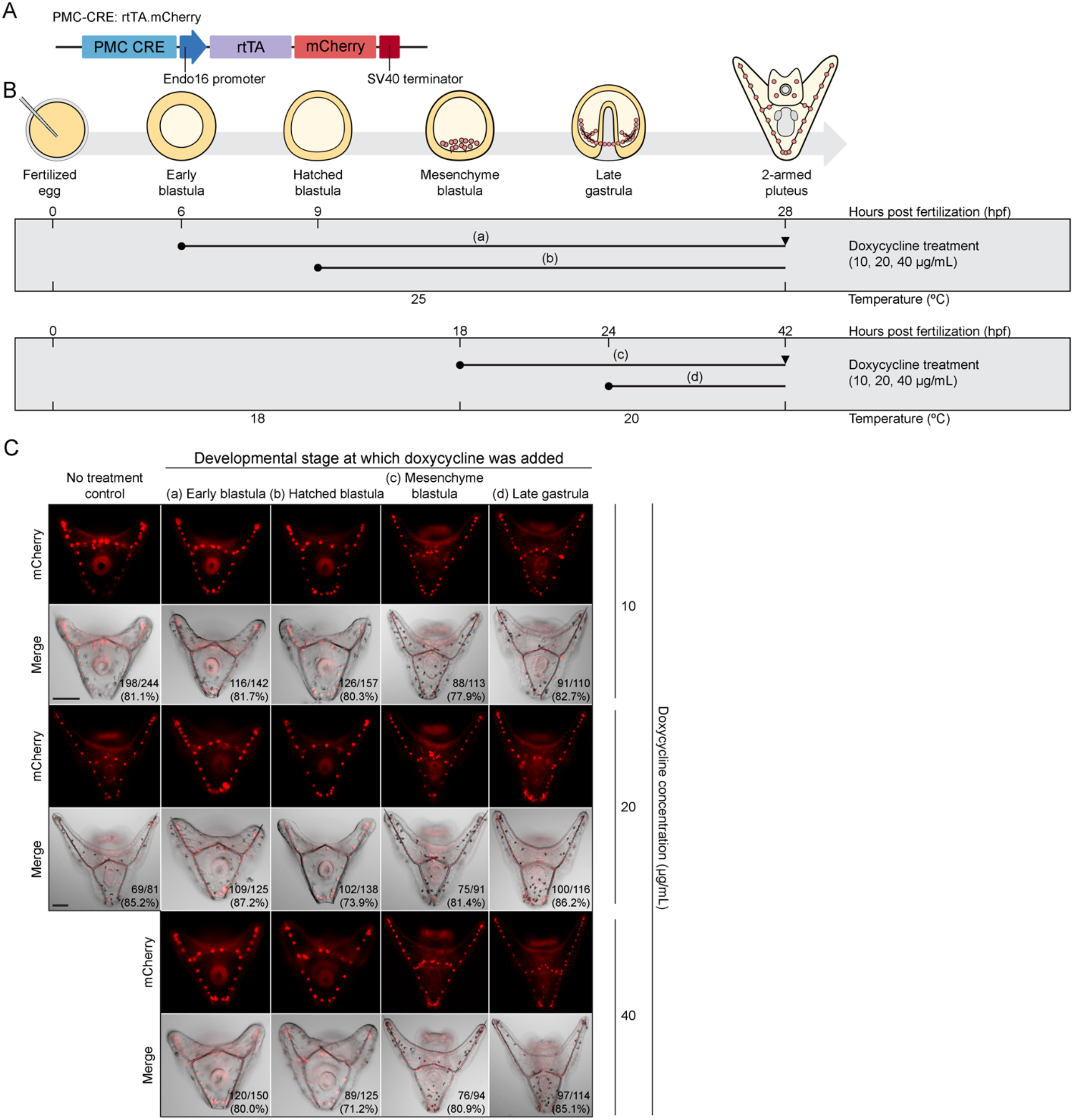
Development of transgenic sea urchin embryos expressing PMC-specific rtTA.mCherry is not affected by doxycycline. (A) Schematic representation of the transgenic activator construct (PMC-CRE: rtTA.mCherry). (B) Experimental design showing the treatment schedule and incubation temperatures. Doxycycline at concentrations of 10, 20, and 40 μg/mL was added to injected embryos at various developmental stages (a-d). Solid circles indicate the stages at which doxycycline was added. Arrowheads indicate the stages at which embryos were collected for analysis. (C) Representative images of transgenic 2-armed plutei treated with doxycycline. The numbers of injected embryos expressing rtTA.mCherry that reached the 2-armed pluteus stage were scored. Top row: mCherry fluorescence in live embryos; Bottom row: mCherry fluorescence overlayed onto DIC images. Scale bar: 50 μm

**Figure S4:**
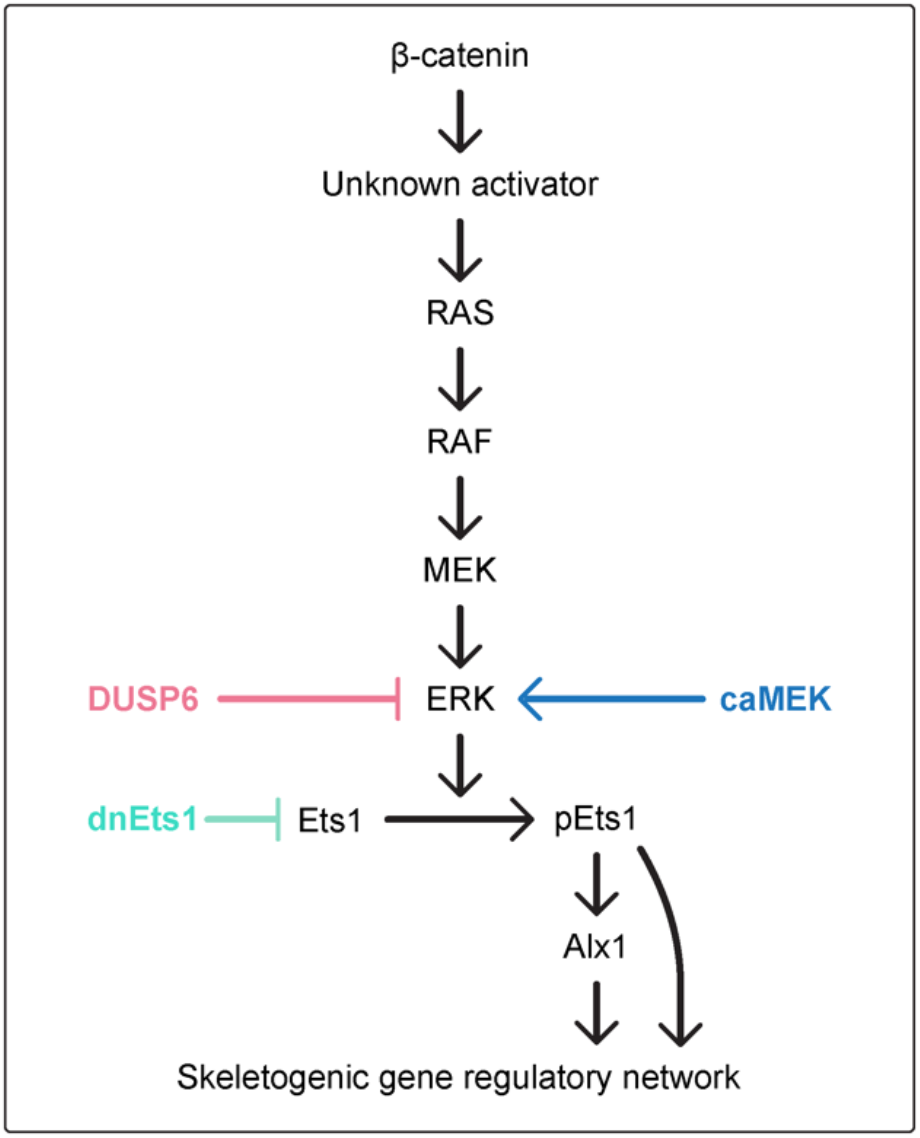
Model for the MAPK signaling pathway involved in regulating sea urchin skeletogenesis (adapted from Röttinger et al., 2004). Genes of interest, (1) dominant negative Ets1 (dnEts1), (2) constitutively active mitogen-activated protein kinase kinase (caMEK), and (3) dual specificity phosphatase 6 (DUSP6) were overexpressed in PMCs using the Tet-On inducible system.

**Figure S5:**
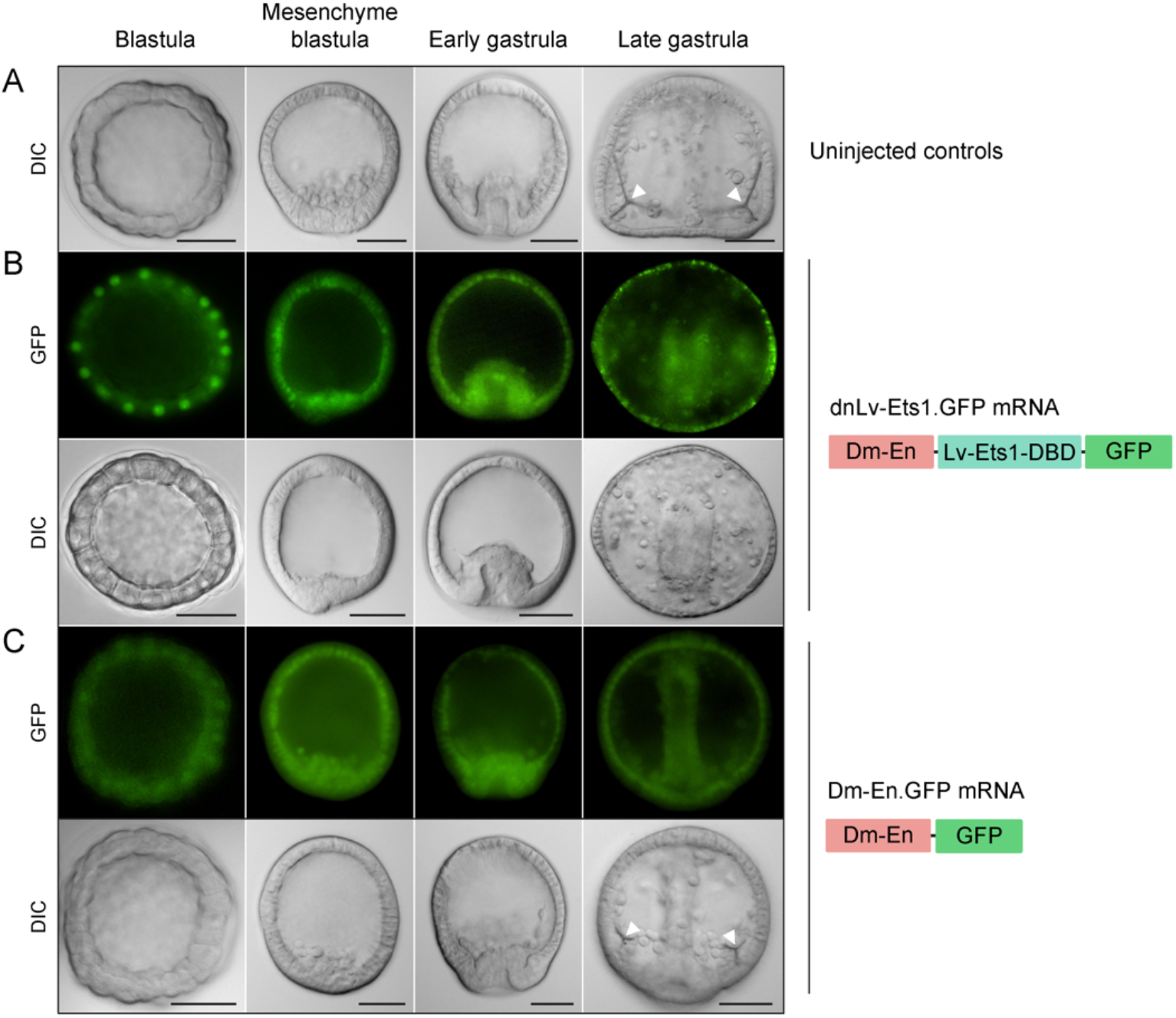
Overexpression of dominant negative Lv-Ets1 (dnLv-Ets1.GFP) inhibits PMC specification and skeletogenesis. (A) DIC images of uninjected control embryos. (B) Embryos injected with capped mRNA encoding dnLv-Ets1.GFP, exhibiting inhibition of PMC ingression and skeletogenesis. Top row: GFP fluorescence in live embryos. Bottom row: GFP fluorescence overlayed onto DIC images. (C) Embryos injected with capped mRNA encoding the *Drosophila* Engrailed repressor domain fused to GFP (Dm-En.GFP) developed normally. Top row: GFP fluorescence in live embryos. Bottom row: GFP fluorescence overlayed onto DIC images. Arrowheads indicate spicule primordia. Scale bar: 50 μm.

**Figure S6:**
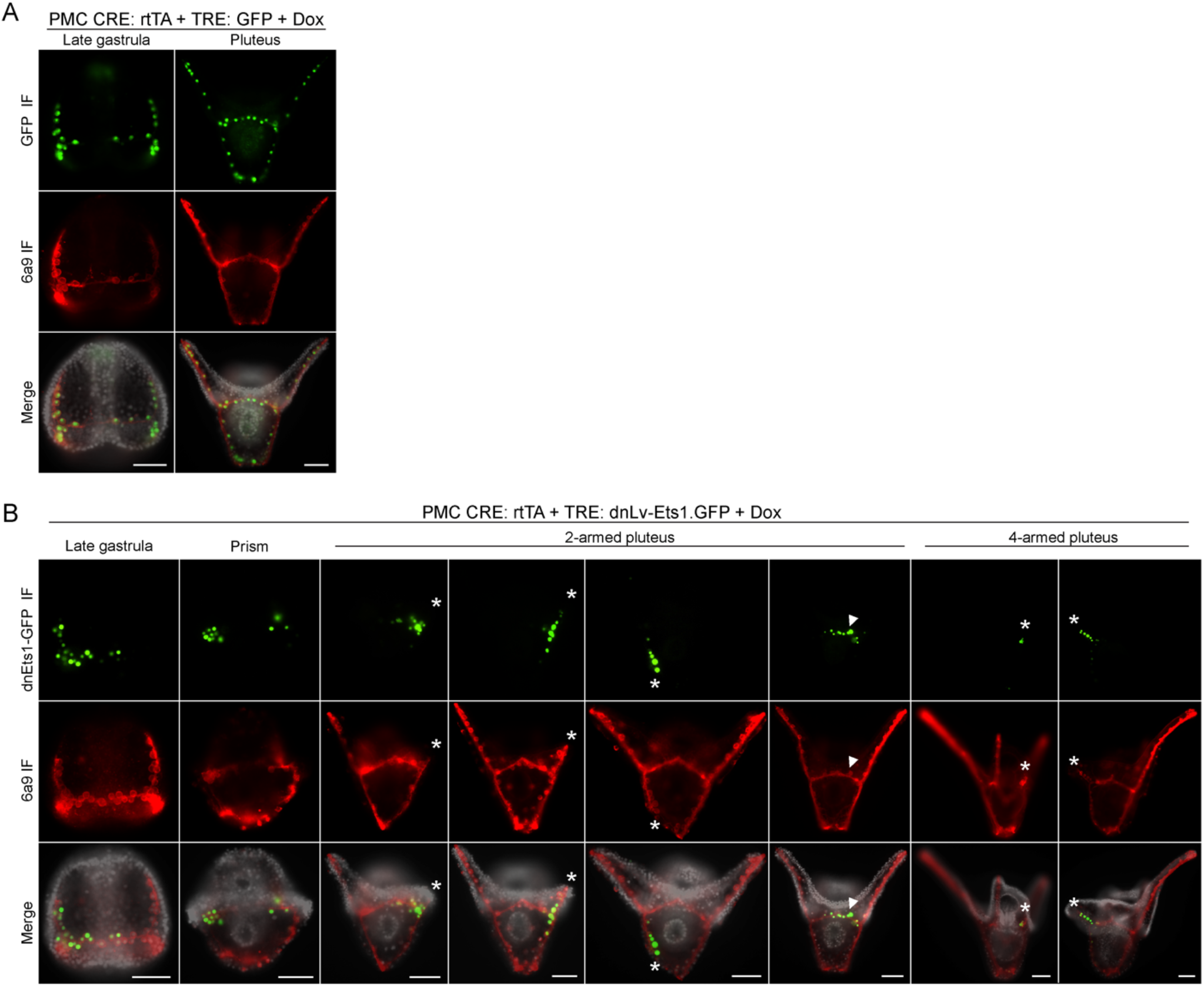
PMC marker (6a9) and GFP immunostaining of transgenic embryos with induced GFP or dnLv-Ets1.GFP expression. (A) Transgenic embryos with induced GFP expression showed normal PMC specification and skeletal growth. Top row: GFP-immunostained cells; Middle row: MSP130-immunostained skeletal structures (6a9 antibody); Bottom row: Fluorescence merged with Hoechst 33342 counterstain in grayscale. (B) Transgenic embryos with asymmetric expression of PMC-specific dnLv-Ets1-GFP show an inhibition of skeletal growth and elongation (asterisks). Expression of dnLv-Ets1.GFP did not affect MSP130 expression. Expression of dnLv-Ets1.GFP in the ventral transverse rods also did not affect skeletogenesis in 2-armed plutei (arrowheads). Top row: GFP-immunostained cells; Middle row: 6a9-immunostained skeletal structures; Bottom row: Fluorescence merged with Hoechst 33342 counterstain in grayscale.

**Figure S7:**
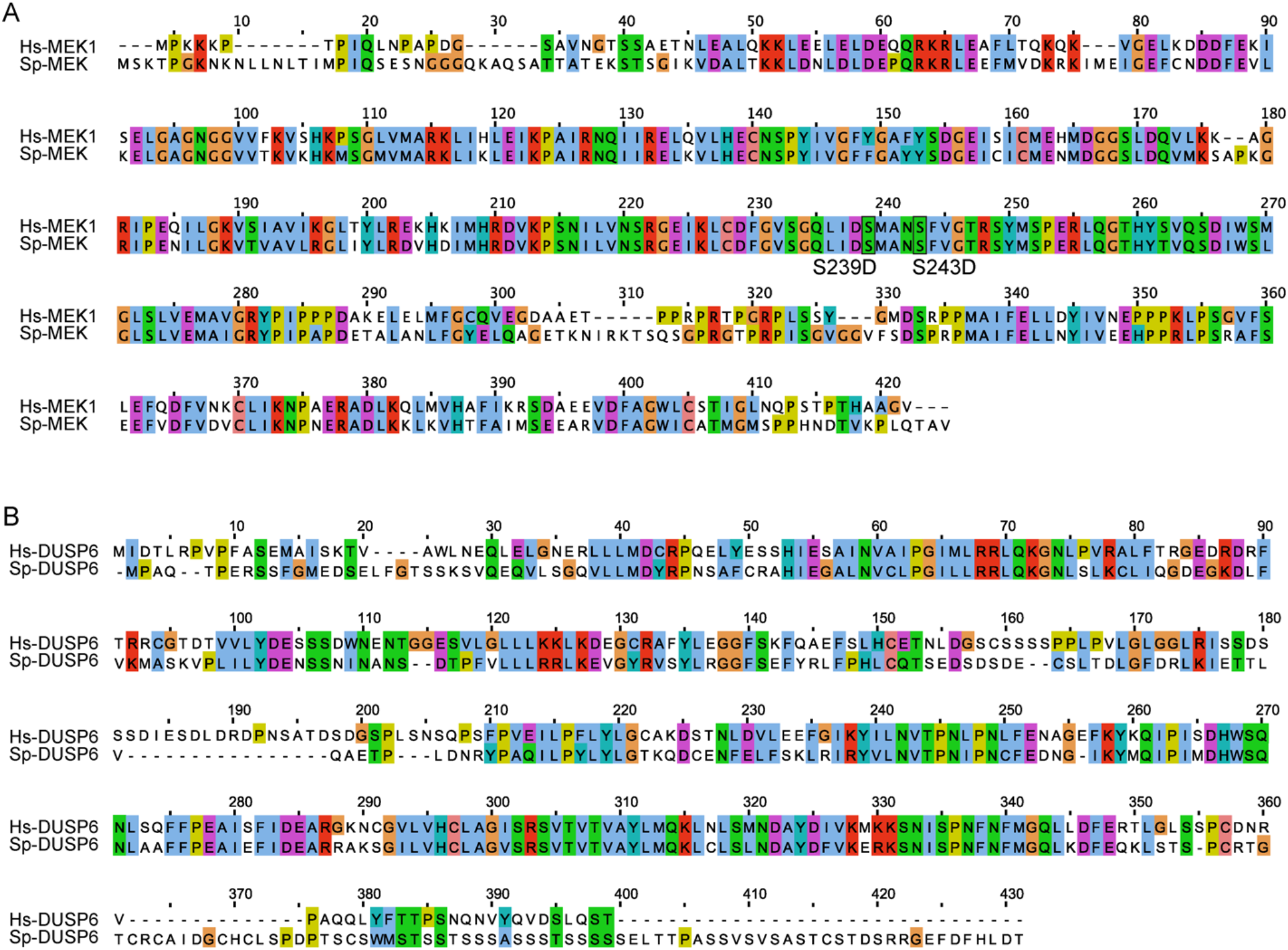
Sequence alignment of *S. purpuratus* and human MEK and DUSP6 proteins. (A) Clustal Omega alignment of *S. purpuratus* MEK (Sp-MEK) and human MEK1 (Hs-MEK1) proteins reveals a high degree of sequence conservation. Mutations that generate constitutively active/phosphomimetic MEK (Sp: S239D/S243D; Hs: S218D/S222D) are shown (Brunet et al., 1994). (B) Clustal Omega alignment of *S. purpuratus* DUSP6 (Sp-DUSP6) and human DUSP6 (Hs-DUSP6) proteins.

**Figure S8:**
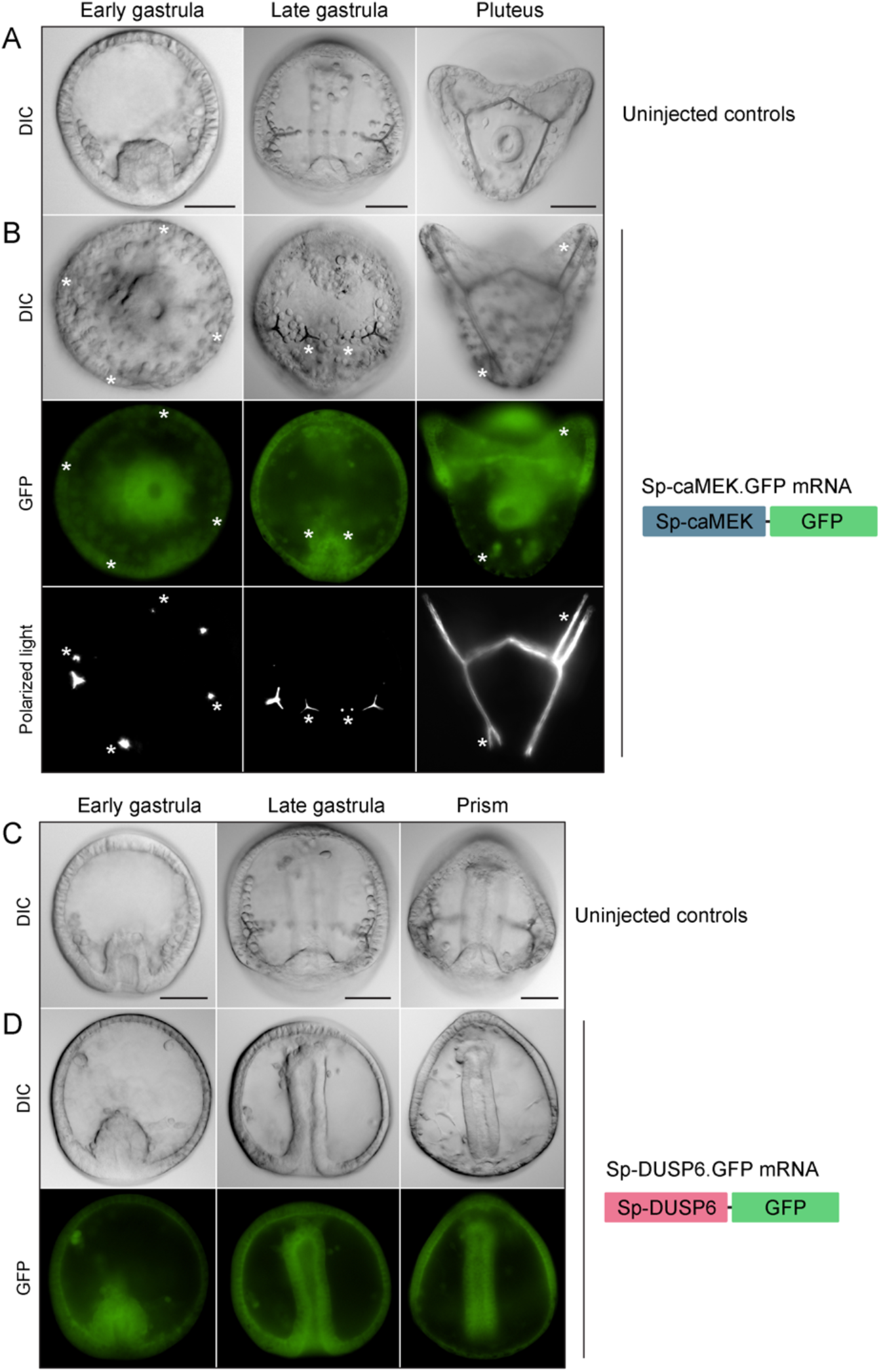
Overexpression of constitutively active *S. purpuratus* MEK (Sp-caMEK.GFP) and DUSP6 (Sp-DUSP6.GFP) disrupts skeletogenesis. (A) DIC images of uninjected control embryos. (B) Overexpression of Sp-caMEK.GFP resulted in supernumerary spicules and abnormal branching (asterisks). Top row: DIC images of injected embryos; Middle row: GFP fluorescence in live embryos; Bottom row: Polarized light images showing skeletal elements. (C) DIC images of uninjected control embryos. (D) Overexpression of Sp-DUSP6.GFP inhibited PMC specification and skeletogenesis. Top row: DIC images of injected embryos; Bottom row: GFP fluorescence in live embryos. Scale bar: 50 μm.

**Supplemental Table S1:** GFP ELISA analysis of transgenic embryos showing Dox dose-and time-dependent induction of gene expression.

## Materials and Methods

### Animals

Adult *Lytechinus variegatus* were acquired from Pelagic Corp (Sugarloaf Key, FL, USA). Adult *Strongylocentrotus purpuratus* were obtained from Pete Halmay (San Diego, CA, USA). Spawning was induced by intracoelomic injection of 0.5 M KCl. *L. variegatus* embryos were cultured in artificial seawater (ASW) at 18-25°C in temperature-controlled incubators while *S. purpuratus* embryos were cultured at 15°C. Feeding-stage *L. variegatus* larvae were fed with *Rhodomonas lens* algae.

### Plasmid constructs

The EpGFPII plasmid (Cameron et al., 2004), which contains the sea urchin-specific *S. purpuratus endo16* (*Sp-endo16*) basal promoter, was used as a backbone vector. Several changes were introduced to the plasmid; most significantly, the *S. purpuratus cyIIa* leader sequence positioned immediately upstream of the GFP coding sequence was removed and the minimal SV40 terminator sequence was replaced with a longer version of the SV40 poly(A) terminator sequence (see supplemental Materials and Methods).

#### Transgenic activator constructs

The reverse tetracycline-controlled transactivator (rtTA) recombinant gene, based on the rtTA-Advanced sequence from pSLIK-Neo (Shin et al., 2006), was synthesized as gBlock gene fragments with and without an mCherry tag by Integrated DNA Technologies (Coralville, IA, USA). The gBlocks were cloned into EpGFPII in place of the GFP coding sequence downstream of the *Sp-endo16* promoter. To drive PMC-specific expression, a *Sp-EMI/TM* intronic *cis*-regulatory element (CRE) (characterized in Khor et al., 2019) was cloned upstream of the promoter to generate PMC-CRE: rtTA and PMC-CRE: rtTA.mCherry. For inducible GFP expression in *S. purpuratus* embryos, a different pair of transactivator and responder were used. The tetON3G recombinant gene, based on the transactivator sequence from pCAG-TetON-3G (Faedo et al., 2017), was synthesized as a gBlock gene fragment by Integrated DNA Technologies. CREs that were shown to drive expression specifically in the pigment cells (*Sp-pks1* and *Sp-fmo2*) (Khor et al., 2021), gut (*Sp-endo16*) (Yuh and Davidson, 1996), and oral ectoderm (*Sp-nodal*) (Nam et al., 2007) were cloned upstream of the tetON3G gene (see supplemental Materials and Methods).

#### Transgenic responder constructs

The tetracycline response element (TRE) and minimal CMV promoter were PCR-amplified from pInducer20 (Meerbrey et al., 2011) and cloned upstream of the GFP coding sequence to generate TRE: GFP. The *Drosophila* Engrailed repressor domain (Dm-En) coding sequence was PCR-amplified from genomic DNA (Margolin et al., 1994). The coding sequences for *L. variegatus* Ets1 DNA binding domain (Lv-Ets1-DBD), constitutively active/phosphomemetic *S. purpuratus* MEK (LOC576066, Sp-caMEK S239D/S243D) (Brunet et al., 1994), *S. purpuratus* dual specificity phosphatase 6 (LOC115919104, Sp-DUSP6), and *Vibrio vulnificus* nitroreductase (NTR-2.0) (Sharrock et al., 2022) were synthesized as gBlock gene fragments with flanking restriction sites by Integrated DNA Technologies (Coralville, IA, USA) and cloned upstream of GFP in the TRE: GFP construct (see supplemental Materials and Methods). For inducible GFP expression in *S. purpuratus* embryos, the TRE3Gp promoter containing the TRE, minimal CMV promoter, and CMV 5’-UTR was cloned upstream of the GFP coding sequence to generate TRE3Gp: GFP (Kang et al., 2019).

### Capped mRNA synthesis

Capped mRNAs were synthesized using the mMessage mMachine SP6 Transcription Kit (Invitrogen/Thermo Fisher Scientific, Waltham, MA, USA). PCR products containing a 5’ SP6 promoter and a 3’ SV40 poly(A) terminator sequence were used as templates for the *in vitro* transcription reactions (see supplemental Materials and Methods).

### Microinjection

Linearized plasmids were injected into fertilized *L. variegatus* eggs following established protocols (Arnone et al., 2004; Cheers and Ettensohn, 2004). Each 20 μL injection solution contained 75 ng of the transactivator plasmid, 75 ng of the responder plasmid, 500 ng of HindIII-digested genomic DNA, 0.12 M KCl, 20% glycerol, and 0.1% Texas Red-Dextran (10,000 MW). Linear DNA injected into fertilized sea urchin eggs forms a large concatemer that is randomly inherited by one or a few cells during cleavage (McMahon et al., 1985). For mRNA overexpression assays, each 5 μL injection solution contained 1.0 μg/μL mRNA, 0.12 M KCl, 20% glycerol, and 0.1% Texas Red-Dextran (10,000 MW).

### Embryo culture drug treatments

A 10 mg/mL stock solution of Doxycycline hyclate (Dox) (D9891, Sigma-Aldrich: St. Louis, MO, USA) was prepared in sterile H_2_O and stored in light-protected microcentrifuge tubes at -20°C. Dox was added directly to cultures at different developmental stages. Metronidazole (MTZ) (M3761, Sigma-Aldrich, St. Louis, MO, USA) was dissolved directly in ASW to obtain a final concentration of 1 mM. To induce nitroreductase-mediated cell ablation, embryos were collected by centrifugation and resuspended in ASW containing 1 mM MTZ and 5 μg/mL doxycycline.

### GFP ELISA

To determine the effect of doxycycline concentration on transgene expression, approximately 3000 eggs were injected with PMC-CRE: rtTA and TRE: GFP and incubated at 20°C overnight. The embryos were then pooled at the late gastrula stage and divided into 8 parallel cultures. GFP expression was induced overnight using a range of doxycycline concentrations (0, 2, 5, 10, 20, 40, 80, and 160 μg/mL). At the 2-armed pluteus stage (approximately 48 hours post fertilization), protein was extracted from the 8 cultures using the lysis buffer from the GFP Fluorescent ELISA Kit (ab229403) (Abcam, Cambridge, UK) that was supplemented with Halt Protease Inhibitor Cocktail (Thermo Fisher Scientific, Waltham, MA, USA). The total protein concentration for each culture was measured using the Pierce BCA Protein Assay Kit (Thermo Fisher Scientific, Waltham, MA, USA). Using the same amount of protein from each sample, GFP concentration was then determined using the GFP Fluorescent ELISA Kit. Relative GFP levels were then calculated using min-max normalization. To determine the earliest time point at which doxycycline induced GFP expression could be detected, approximately 3000 eggs were injected and incubated at 23°C overnight. The embryos were then pooled at the late gastrula stage and divided into 8 cultures. GFP expression was induced via the addition of 5 μg/mL doxycycline. Total protein was then extracted every hour (0, 1, 2, 3, 4, 5, 6, and 7 hours post doxycycline treatment) and GFP levels were measured using the GFP Fluorescent ELISA Kit as described above.

### Combined whole-mount fluorescent in situ hybridization and immunostaining (ImmunoFISH)

ImmunoFISH was employed for simultaneous visualization of *p16* or *sm30b* transcripts and GFP protein. DNA templates for RNA probe synthesis were PCR-amplified with reverse primers that contained a T3 promoter sequence (see supplemental Materials and Methods). Digoxigenin-labeled RNA probes were synthesized using the MEGAscript T3 Transcription Kit (Invitrogen/Thermo Fisher Scientific, Waltham, MA, USA). The single-color fluorescent in situ hybridization (FISH) portion of the protocol was carried out as previously described (Ettensohn et al., 2007; Sharma et al., 2010) with a few modifications. Embryos were collected and fixed at the desired stage for 1 hour in 4% paraformaldehyde (PFA) in ASW followed by at least 15 minutes incubation in ice-cold 100% methanol. Fixed embryos were processed immediately or stored in 100% methanol. After overnight probe hybridization at 55°C, the samples were washed with phosphate buffered saline containing 0.1% Tween-20 (PBST) and transferred to round-bottom 96-well plates. The samples were then blocked for 1 hour at room temperature in PBS containing 4% goat serum, 4% sheep serum, and 1% BSA. This was followed by a 2-hour incubation in an antibody mixture composed of a 1:1000 dilution of anti-digoxigenin-POD (Roche/Sigma-Aldrich, St. Louis, MO, USA) and a 1:500 dilution of anti-GFP antibody (ab6556) (Abcam, Cambridge, UK) in blocking buffer at room temperature. After several washes with PBST, the samples were incubated in 1:500 DyLight 488 anti-rabbit IgG (Jackson Immuno Research Labs, West Grove, PA, USA) for 2 hours at room temperature. Following several additional PBST washes, the samples were incubated in a 1:100 dilution of Cy3-Tyramide Signal Amplification Solution (TSA plus Cyanine 3 Kit, Akoya Biosciences, Marlborough, MA, USA) for 5 minutes at room temperature. The samples were then counterstained with 0.5 μg/mL Hoechst 33342 in PBST for 12 minutes. Finally, the samples were mounted on slides in anti-fade solution (2.5% DABCO, 50% glycerol, 50% PBS) for examination.

### Immunostaining

Immunstaining of fixed embryos were carried out as previously described (Khor et al., 2017). PMCs were immunostained with monoclonal antibody (mAb) 6a9, which recognizes PMC-specific cell surface proteins of the MSP130 family (Ettensohn and McClay, 1988; Illies et al., 2002). Embryos were fixed at the desired stage with 2% PFA in ASW and transferred to round-bottom 96-well plates for further processing. Fixed embryos expressing GFP were incubated in blocking buffer (5% goat serum and 1% BSA in PBS) containing a 1:2 dilution of 6a9 tissue culture supernatant and a 1:500 dilution of the anti-GFP antibody. Following several washes with PBST, they were incubated in blocking buffer containing a 1:100 dilution of Alexa Fluor 594 anti-rabbit IgG and a 1:500 dilution of DyLight 488 anti-rabbit IgG (Jackson Immuno Research Labs, West Grove, PA, USA) for 2 hours at room temperature. They were counterstained with Hoechst 33342 (0.5 μg/mL) in PBST for 12 minutes and mounted on slides in anti-fade solution for examination.

## Supplemental Methods and Materials

### Plasmid construction

>Sp_endo16_promoter

TTAAACTGTTTGAGTTTCGTCTCCTGATTGTGCTATCAAAGACAAAGGGGTGTAACTTTACCCCCCTCATCAAGAGCGGAGGGTTAAAtAGAGAAAgACTGGTcgAGGACAgGTCATAATATTGCTAATTTTTAAGCTTATCATC

>SV40_PA_Terminator

TGATCATAATCAAGCCATATCACATCTGTAGAGGTTTACTTGCTTTAAAAAACCTCCACACCTCCCCCTGAACCTGAAACATAAAATGAATGCAATTGTTGTTGTTAACTTGTTTATTGCAGCTTATAATGGTTACA<u>AATAAA</u>GCAATAGCATCACAAATTTCACAAATAAAGCATTTTTTTCACTGCATTCTAGTTGTGGTTTGTCCAAACTCATCAATGTATCTTATCATGTCTGGATCTGC

>rtTA

ATGTCTAGACTGGACAAGAGCAAAGTCATAAACGGAGCTCTGGAATTACTCAATGGTGTCGGTATCGAAGGCCTGACGACAAGGAAACTCGCTCAAAAGCTGGGAGTTGAGCAGCCTACCCTGTACTGGCACGTGAAGAACAAGCGGGCCCTGCTCGATGCCCTGCCAATCGAGATGCTGGACAGGCATCATACCCACTTCTGCCCCCTGGAAGGCGAGTCATGGCAAGACTTTCTGCGGAACAACGCCAAGTCATACCGCTGTGCTCTCCTCTCACATCGCGACGGGGCTAAAGTGCATCTCGGCACCCGCCCAACAGAGAAACAGTACGAAACCCTGGAAAATCAGCTCGCGTTCCTGTGTCAGCAAGGCTTCTCCCTGGAGAACGCACTGTACGCTCTGTCCGCCGTGGGCCACTTTACACTGGGCTGCGTATTGGAGGAACAGGAGCATCAAGTAGCAAAAGAGGAAAGAGAGACACCTACCACCGATTCTATGCCCCCACTTCTGAGACAAGCAATTGAGCTGTTCGACCGGCAGGGAGCCGAACCTGCCTTCCTTTTCGGCCTGGAACTAATCATATGTGGCCTGGAGAAACAGCTAAAGTGCGAAAGCGGCGGGCCGACCGACGCCCTTGACGATTTTGACTTAGACATGCTCCCAGCCGATGCCCTTGACGACTTTGACCTTGATATGCTGCCTGCTGACGCTCTTGACGATTTTGACCTTGACATGCTCCCCGGGTGA

>rtTA_mCherry

ATGTCTAGACTGGACAAGAGCAAAGTCATAAACGGAGCTCTGGAATTACTCAATGGTGTCGGTATCGAAGGCCTGACGACAAGGAAACTCGCTCAAAAGCTGGGAGTTGAGCAGCCTACCCTGTACTGGCACGTGAAGAACAAGCGGGCCCTGCTCGATGCCCTGCCAATCGAGATGCTGGACAGGCATCATACCCACTTCTGCCCCCTGGAAGGCGAGTCATGGCAAGACTTTCTGCGGAACAACGCCAAGTCATACCGCTGTGCTCTCCTCTCACATCGCGACGGGGCTAAAGTGCATCTCGGCACCCGCCCAACAGAGAAACAGTACGAAACCCTGGAAAATCAGCTCGCGTTCCTGTGTCAGCAAGGCTTCTCCCTGGAGAACGCACTGTACGCTCTGTCCGCCGTGGGCCACTTTACACTGGGCTGCGTATTGGAGGAACAGGAGCATCAAGTAGCAAAAGAGGAAAGAGAGACACCTACCACCGATTCTATGCCCCCACTTCTGAGACAAGCAATTGAGCTGTTCGACCGGCAGGGAGCCGAACCTGCCTTCCTTTTCGGCCTGGAACTAATCATATGTGGCCTGGAGAAACAGCTAAAGTGCGAAAGCGGCGGGCCGACCGACGCCCTTGACGATTTTGACTTAGACATGCTCCCAGCCGATGCCCTTGACGACTTTGACCTTGATATGCTGCCTGCTGACGCTCTTGACGATTTTGACCTTGACATGCTCCCCGGGGGAGGTGGAGGTTCAGGTGGAGGAGGTTCCATGGTGAGCAAGGGCGAGGAGGATAACATGGCCATCATCAAGGAGTTCATGCGCTTCAAGGTGCACATGGAGGGCTCCGTGAACGGCCACGAGTTCGAGATCGAGGGCGAGGGCGAGGGCCGCCCCTACGAGGGCACCCAGACCGCCAAGCTGAAGGTGACCAAGGGTGGCCCCCTGCCCTTCGCCTGGGACATCCTGTCCCCTCAGTTCATGTACGGCTCCAAGGCCTACGTGAAGCACCCCGCCGACATCCCCGACTACTTGAAGCTGTCCTTCCCCGAGGGCTTCAAGTGGGAGCGCGTGATGAACTTCGAGGACGGCGGCGTGGTGACCGTGACCCAGGACTCCTCCCTGCAGGACGGCGAGTTCATCTACAAGGTGAAGCTGCGCGGCACCAACTTCCCCTCCGACGGCCCCGTAATGCAGAAGAAGACCATGGGCTGGGAGGCCTCCTCCGAGCGGATGTACCCCGAGGACGGCGCCCTGAAGGGCGAGATCAAGCAGAGGCTGAAGCTGAAGGACGGCGGCCACTACGACGCTGAGGTCAAGACCACCTACAAGGCCAAGAAGCCCGTGCAGCTGCCCGGCGCCTACAACGTCAACATCAAGTTGGACATCACCTCCCACAACGAGGACTACACCATCGTGGAACAGTACGAACGCGCCGAGGGCCGCCACTCCACCGGCGGCATGGACGAGCTGTACAAGTGA

>SpEMI_CRE

CCCTATAATAACGGTATGGTTTTAATACTGTAATGATTTGTCCAAATACCATTGTCTAAATATCAAAGGCCACATACCAGACACTTAATAATATGATATACCCGAAGGAGTCTGTGGTTGAACCTGAGCTTCGAAGCGAAATATTGTGATTGCATAACATCCTTCTCATGTTTTGACATACAGTCATAAATTAAACCGTTAATGAAAACATTTCAAAACAAACTTCCCCTCCAGCATGGCCTTAAACCATCCTTCTCAGAGGAATAAATTCATTTTTCTCCAAAATAGGAATTAGAGATTCCCTTGAATGCTGCGGCAACGCAGGGGATGTTGGCTAATTGACATAATTTAGCAATTATGTTGAACTGCCACGGTACAAGTATATATTCTCGTGCGAAACAACTAAGTGCAATAAGGCGTTTGTTATTTCGTCGGGGCGTTAATAGATTTAAACTTTTTCGTCCGCCTGTAAAGTGATAACAAAATTAACAGGTAGACAACATACTCCGAGAGACTAACCGTCATTTCAAGAGAATGATTCAAGAAGTTATTATCACCATGCACATGAATAATTTTAGCTGATAAGTGAAAGAAGAAAGTGATAATTAATGTGGTGTCACTTCCTCGATTTTGACGTTGTGCTTGACGGAAATATGGAAGGCATTGGTTGATAAACCAACCACAACGAGATGAAGATAGTAATCTTATGTCTGGCTGAGTTTAGTGCAATTTGTTTCATTGAAAAGATGAGCAGCCACTTGACGTGCATTTAAGAATACCTTGTGTAAATAATTAGGCC

>TRE_mCMV

TTTACCACTCCCTATCAGTGATAGAGAAAAGTGAAAGTCGAGTTTACCACTCCCTATCAGTGATAGAGAAAAGTGAAAGTCGAGTTTACCACTCCCTATCAGTGATAGAGAAAAGTGAAAGTCGAGTTTACCACTCCCTATCAGTGATAGAGAAAAGTGAAAGTCGAGTTTACCACTCCCTATCAGTGATAGAGAAAAGTGAAAGTCGAGTTTACCACTCCCTATCAGTGATAGAGAAAAGTGAAAGTCGAGTTTACCACTCCCTATCAGTGATAGAGAAAAGTGAAAGTCGAGCTCGGTACCCGGGTCGAGGTAGGCGTGTACGGTGGGAGGCCTATATAAGCAGAGCTCGTTTAGTGAACCGTCAGATC

>GFP

ATGAGTAAAGGAGAAGAACTTTTCACTGGAGTTGTCCCAATTCTTGTTGAATTAGATGGTGATGTTAATGGGCACAAATTTTCTGTCAGTGGAGAGGGTGAAGGTGATGCAACATACGGAAAACTTACCCTTAAATTTATTTGCACTACTGGAAAACTACCTGTTCCATGGCCAACACTTGTCACTACTCTCACTTATGGTGTTCAATGCTTTTCAAGATACCCAGATCATATGAAACGGCATGACTTTTTCAAGAGTGCCATGCCCGAAGGTTATGTACAGGAAAGAACTATATTTTTCAAAGATGACGGGAACTACAAGACACGTGCTGAAGTCAAGTTTGAAGGTGATACCCTTGTTAATAGAATCGAGTTAAAAGGTATTGATTTTAAAGAAGATGGAAACATTCTTGGACACAAATTGGAATACAACTATAACTCACACAATGTATACATCATGGCAGACAAACAAAAGAATGGAATCAAAGTTAACTTCAAAATTAGACACAACATTGAAGATGGAAGCGTTCAACTAGCAGACCATTATCAACAAAATACTCCAATTGGCGATGGCCCTGTCCTTTTACCAGACAACCATTACCTGTCCACACAATCTGCCCTTTCGAAAGATCCCAACGAAAAGAGAGACCACATGGTCCTTCTTGAGTTTGTAACAGCTGCTGGGATTACACATGGCATGGATGAACTATACAAATAA

>DmEn_LvEts1DBD_GFP

ATGGCCCTGGAGGATCGCTGCAGTCCACAGTCAGCGCCCAGCCCCATTACCCTACAAATGCAGCATCTTCACCACCAGCAACAGCAGCAGCAGCAACAGCAGCAGCAAATGCAGCACCTCCACCAGCTGCAGCAACTGCAGCAGTTGCACCAACAGCAACTGGCCGCCGGTGTCTTCCACCATCCGGCAATGGCCTTCGATGCCGCTGCAGCCGCCGCTGCTGCAGCTGCTGCTGCGGCCGCCCACGCTCATGCTGCTGCACTGCAGCAGCGCCTCAGTGGCAGTGGATCGCCCGCATCCTGCTCCACGCCCGCCTCGTCCACGCCGCTGACCATCAAGGAGGAGGAAAGCGACTCCGTGATCGGTGACATGAGTTTCCACAATCAGACGCACACCACCAACGAGGAGGAGGAGGCGGAGGAGGATGACGACATTGATGTGGATGTGGATGATACGTCGGCGGGCGGACGCCTGCCACCACCCGCCCACCAGCAGCAGTCGACGGCCAAGCCCTCGCTGGCCTTTTCCATCTCCAACATCCTGAGCGATCGTTTCGGAGATGTCCAGAAGCCGGGCAAGTCGATGGAGAACCAGGCCAGCATATTCCGCCCCTTCGAGGCGAGTCGTTCCCAGACTGCCACGCCCTCCGCCTTTACAAGAGTGGATCTGCTGGAGTTTAGCCGGCAACAGCAGGCTGCCGCCGCAGCCGCTACTGCGGCCATGATGCTGGAACGGGCCAACTTCCTTAACTGCTTCAATCCGGCTGCCTATCCCAGGATACACGAGGAAATCGTGCAGAGTCGGCTGCGCAGGAGTGCAGCCAATGCCGTCATCCCGCCGCCCATGAGCTCCAAGATGAGCGATGCCAATCCAGAGAAATCTGCTCTGGGATCCGGAGGTGGAGGTTCAGCTAGCGGTGGAGGAGGTTCCATGTCTGACTCACCCCTCGCAGGCGATGTGGTATCAAACCCCGTTATCCCCGCGGCCGTACTAGCTGGTTATTCCGGAAGCGGACCCATTCAACTCTGGCAATTTCTCCTGGAGCTTCTGACCGACAAGACTTGCCAGCACATCATCAGTTGGACGGGCGATGGCTGGGAATTCAAGTTGTCCGACCCCGATGAGGTTGCCCGACGATGGGGCAAGCGCAAGAACAAGCCCAAGATGAACTATGAGAAGCTGAGCCGCGGACTGCGCTATTATTACGACAAGAACATCATCCACAAGACGGCGGGCAAGCGCTATGTCTACCGCTTCGTCTGCGACCTGCAGAGCCTGCTGGGATACTCGCCCGAGGAGCTGCACGAGATGGTCGGCGTCAGCCCCGCGCGAGACGACGACGGTGGTGGAGGATCGGGAGGAGGTGGTTCTTCTAGAATGAGTAAAGGAGAAGAACTTTTCACTGGAGTTGTCCCAATTCTTGTTGAATTAGATGGTGATGTTAATGGGCACAAATTTTCTGTCAGTGGAGAGGGTGAAGGTGATGCAACATACGGAAAACTTACCCTTAAATTTATTTGCACTACTGGAAAACTACCTGTTCCATGGCCAACACTTGTCACTACTCTCACTTATGGTGTTCAATGCTTTTCAAGATACCCAGATCATATGAAACGGCATGACTTTTTCAAGAGTGCCATGCCCGAAGGTTATGTACAGGAAAGAACTATATTTTTCAAAGATGACGGGAACTACAAGACACGTGCTGAAGTCAAGTTTGAAGGTGATACCCTTGTTAATAGAATCGAGTTAAAAGGTATTGATTTTAAAGAAGATGGAAACATTCTTGGACACAAATTGGAATACAACTATAACTCACACAATGTATACATCATGGCAGACAAACAAAAGAATGGAATCAAAGTTAACTTCAAAATTAGACACAACATTGAAGATGGAAGCGTTCAACTAGCAGACCATTATCAACAAAATACTCCAATTGGCGATGGCCCTGTCCTTTTACCAGACAACCATTACCTGTCCACACAATCTGCCCTTTCGAAAGATCCCAACGAAAAGAGAGACCACATGGTCCTTCTTGAGTTTGTAACAGCTGCTGGGATTACACATGGCATGGATGAACTATACAAATAA

>Sp_caMEK_GFP

ATGTCGAAGACGCCCGGAAAGAACAAAAATTTATTGAACTTGACAATAATGCCAATACAATCGGAATCAAATGGCGGAGGACAGAAAGCACAATCTGCAACTACAGCGACAGAGAAAAGCACCTCTGGAATTAAAGTGGATGCACTGACAAAGAAACTGGACAATCTAGATCTTGATGAACCCCAAAGGAAACGTCTTGAAGAATTTATGGTGGACAAGAGGAAGATCATGGAAATTGGAGAATTCTGCAATGATGACTTTGAAGTGCTTAAGGAATTAGGAGCGGGGAATGGAGGCGTGGTCACTAAAGTGAAACATAAGATGAGCGGTATGGTAATGGCAAGAAAGCTCATTAAACTTGAAATCAAACCAGCCATCAGAAATCAGATCATCCGAGAACTGAAAGTGCTTCATGAGTGCAATTCACCATACATCGTGGGTTTCTTTGGAGCGTATTACAGCGATGGAGAGATTTGTATCTGCATGGAAAACATGGATGGCGGTTCTTTAGATCAAGTCATGAAGTCAGCACCCAAGGGAAGAATACCAGAAAATATCCTTGGCAAGGTCACAGTTGCAGTTTTGAGGGGGCTCATTTATTTAAGAGATGTGCATGACATAATGCACAGAGATGTGAAACCATCAAATATCCTGGTCAACTCACGGGGAGAGATCAAGTTGTGTGATTTTGGTGTGAGCGGTCAGCTTATCGACGATATGGCAAATGACTTTGTGGGAACAAGGTCATATATGTCGCCTGAGAGACTGCAAGGCACACACTACACAGTACAGTCTGACATCTGGAGTCTTGGCCTATCGCTGGTCGAGATGGCCATTGGACGTTACCCCATCCCCGCCCCTGACGAGACGGCGCTAGCAAACCTCTTCGGCTACGAGCTCCAGGCGGGAGAGACCAAGAATATCCGGAAAACAAGTCAGTCGGGACCTCGCGGTACTCCAAGACCGATCAGTGGAGTTGGAGGAGTGTTTTCAGATTCCCCAAGACCTATGGCCATCTTTGAATTACTCAATTATATTGTAGAAGAGCATCCACCAAGATTACCATCCAGAGCCTTCTCAGAAGAATTTGTAGATTTTGTTGACGTATGTTTGATCAAGAATCCAAATGAGAGGGCAGATCTTAAGAAACTCAAGGTACATACATTTGCAATAATGTCTGAGGAAGCGAGAGTAGACTTTGCTGGATGGATTTGTGCAACCATGGGGATGAGTCCTCCCCATAACGACACGGTCAAACCCCTACAGACAGCTGTCCTCGAGTCTAGAATGAGTAAAGGAGAAGAACTTTTCACTGGAGTTGTCCCAATTCTTGTTGAATTAGATGGTGATGTTAATGGGCACAAATTTTCTGTCAGTGGAGAGGGTGAAGGTGATGCAACATACGGAAAACTTACCCTTAAATTTATTTGCACTACTGGAAAACTACCTGTTCCATGGCCAACACTTGTCACTACTCTCACTTATGGTGTTCAATGCTTTTCAAGATACCCAGATCATATGAAACGGCATGACTTTTTCAAGAGTGCCATGCCCGAAGGTTATGTACAGGAAAGAACTATATTTTTCAAAGATGACGGGAACTACAAGACACGTGCTGAAGTCAAGTTTGAAGGTGATACCCTTGTTAATAGAATCGAGTTAAAAGGTATTGATTTTAAAGAAGATGGAAACATTCTTGGACACAAATTGGAATACAACTATAACTCACACAATGTATACATCATGGCAGACAAACAAAAGAATGGAATCAAAGTTAACTTCAAAATTAGACACAATATTGAAGATGGAAGCGTTCAACTAGCAGACCATTATCAACAAAATACTCCAATTGGCGATGGCCCTGTCCTTTTACCAGACAACCATTACCTGTCCACACAATCTGCCCTTTCGAAAGATCCCAACGAAAAGAGAGACCACATGGTCCTTCTTGAGTTTGTAACAGCTGCTGGGATTACACATGGCATGGATGAACTATACAAATAA

>Sp_DUSP6_GFP

ATGCCCGCTCAGACTCCTGAGAGGTCGTCCTTCGGAATGGAAGACTCAGAACTTTTCGGGACCAGTTCCAAGTCGGTACAGGAGCAAGTTTTGTCCGGCCAAGTTCTCCTCATGGACTACAGACCTAACTCGGCCTTCTGTCGGGCGCATATAGAAGGAGCACTGAACGTTTGCTTACCGGGAATTCTTCTTCGGAGATTACAGAAGGGGAACCTGTCGTTGAAATGTCTGATTCAAGGGGATGAAGGAAAGGACCTTTTCGTGAAAATGGCCAGCAAAGTTCCTCTCATTTTATATGATGAGAACTCGAGCAACATCAATGCAAACTCAGATACTCCATTCGTCCTATTGCTGAGGAGACTCAAAGAAGTTGGTTATCGGGTCAGCTACTTAAGAGGTGGCTTTAGCGAATTCTACAGACTGTTCCCTCACTTATGCCAAACTTCAGAAGACAGCGATTCCGACGAGTGTTCTCTCACAGATCTGGGCTTCGACAGGCTCAAGATCGAGACGACGTTGGTGCAAGCAGAAACCCCATTGGACAACCGATACCCCGCCCAGATCCTGCCCTACCTGTACCTCGGAACCAAGCAGGATTGTGAAAACTTTGAACTCTTCTCCAAACTGAGAATACGCTATGTTCTGAATGTTACGCCCAACATCCCAAACTGCTTTGAGGACAATGGGATCAAATACATGCAGATTCCTATCATGGACCATTGGAGTCAGAATCTGGCAGCTTTCTTTCCTGAAGCCATCGAGTTTATTGACGAAGCTCGCCGTGCCAAGTCTGGTATCCTTGTCCACTGTCTGGCTGGTGTGAGCCGTTCTGTGACTGTAACCGTGGCATACCTCATGCAGAAGCTCTGCCTCTCTCTCAACGACGCGTACGACTTTGTCAAGGAACGCAAGTCCAACATCTCTCCCAACTTCAACTTCATGGGCCAGCTGAAAGACTTTGAGCAGAAGCTTTCGACGAGTCCCTGTCGGACTGGCACATGTCGGTGTGCCATCGATGGCTGCCACTGTCTCAGCCCAGACCCCACCTCATGCTCATGGATGTCGACATCGTCCACGTCCTCGTCGGCGTCCTCATCAACCTCATCATCGTCCAGTGAGCTTACAACTCCAGCCAGCAGTGTCAGTGTCAGCGCCAGCACGTGTTCAACAGACAGTCGGAGAGGGGAGTTTGATTTCCATTTAGATACCACTAGTAGATCTGCTAGCCTCGAGTCTAGAATGAGTAAAGGAGAAGAACTTTTCACTGGAGTTGTCCCAATTCTTGTTGAATTAGATGGTGATGTTAATGGGCACAAATTTTCTGTCAGTGGAGAGGGTGAAGGTGATGCAACATACGGAAAACTTACCCTTAAATTTATTTGCACTACTGGAAAACTACCTGTTCCATGGCCAACACTTGTCACTACTCTCACTTATGGTGTTCAATGCTTTTCAAGATACCCAGATCATATGAAACGGCATGACTTTTTCAAGAGTGCCATGCCCGAAGGTTATGTACAGGAAAGAACTATATTTTTCAAAGATGACGGGAACTACAAGACACGTGCTGAAGTCAAGTTTGAAGGTGATACCCTTGTTAATAGAATCGAGTTAAAAGGTATTGATTTTAAAGAAGATGGAAACATTCTTGGACACAAATTGGAATACAACTATAACTCACACAATGTATACATCATGGCAGACAAACAAAAGAATGGAATCAAAGTTAACTTCAAAATTAGACACAACATTGAAGATGGAAGCGTTCAACTAGCAGACCATTATCAACAAAATACTCCAATTGGCGATGGCCCTGTCCTTTTACCAGACAACCATTACCTGTCCACACAATCTGCCCTTTCGAAAGATCCCAACGAAAAGAGAGACCACATGGTCCTTCTTGAGTTTGTAACAGCTGCTGGGATTACACATGGCATGGATGAACTATACAAATAA

>NTR2_GFP

ATGACTATTGTTCAAGCTGCCCAATCCCGCTACTCCACCAAAGCCTTTGATGCTTCGCGCAAATTGCCTGAAGAGAAAGTCGCGGCAGTGAAAGAGTTAATCCGCATGAGTGCGTCCAGTGTCAACTCGCAACCTTGGCATTTTATTGTCGCGAGCAGTGAAGAGGGAAAAGCGCGCATCGCCAAAGCAACACAAGGTGGTTTTGCTGCTAATGAGCGCAAGATTTTGGATGCTTCTCACGTTGTGGTGTTCTGCGCCAAAACGGCGATTGATGAAGCGTACCTACTCGACCTTTTGGAAAGCGAAGACAAAGATGGCCGCTACGCCGATGTCGAAGCAAAAAATGGCATGCACGCTGGTCGTTCATTTTTCGTCAACATGCACCGCTTTGACTTGAAAGACGCGCATCACTGGATGGAAAAGCAAGTTTACCTCAATGTGGGGACGCTGCTATTGGGTGCTTCTGCGATGGAGATCGACGCGGTGCCAATTGAAGGCTTCGATGCCAAGGTGCTTGATGAGGAGTTTGGTCTGCGTGAGAAGGGCTTTACCAGCGTGGTGATTGTGCCGCTGGGTTACCATAGCGAAGACGATTTTAATGCTAAGCTGCCAAAATCACGTTGGTCAGCAGAGACTGTTTTTACCGAAATCGGAGGTGGAGGTTCAGGTGGAGGAGGTTCCCTCGAGTCTAGAATGAGTAAAGGAGAAGAACTTTTCACTGGAGTTGTCCCAATTCTTGTTGAATTAGATGGTGATGTTAATGGGCACAAATTTTCTGTCAGTGGAGAGGGTGAAGGTGATGCAACATACGGAAAACTTACCCTTAAATTTATTTGCACTACTGGAAAACTACCTGTTCCATGGCCAACACTTGTCACTACTCTCACTTATGGTGTTCAATGCTTTTCAAGATACCCAGATCATATGAAACGGCATGACTTTTTCAAGAGTGCCATGCCCGAAGGTTATGTACAGGAAAGAACTATATTTTTCAAAGATGACGGGAACTACAAGACACGTGCTGAAGTCAAGTTTGAAGGTGATACCCTTGTTAATAGAATCGAGTTAAAAGGTATTGATTTTAAAGAAGATGGAAACATTCTTGGACACAAATTGGAATACAACTATAACTCACACAATGTATACATCATGGCAGACAAACAAAAGAATGGAATCAAAGTTAACTTCAAAATTAGACACAACATTGAAGATGGAAGCGTTCAACTAGCAGACCATTATCAACAAAATACTCCAATTGGCGATGGCCCTGTCCTTTTACCAGACAACCATTACCTGTCCACACAATCTGCCCTTTCGAAAGATCCCAACGAAAAGAGAGACCACATGGTCCTTCTTGAGTTTGTAACAGCTGCTGGGATTACACATGGCATGGATGAACTATACAAATAA

>tetON3G

ATGTCTAGACTGGACAAGAGCAAAGTCATAAACTCTGCTCTGGAATTACTCAATGGAGTCGGTATCGAAGGCCTGACGACAAGGAAACTCGCTCAAAAGCTGGGAGTTGAGCAGCCTACCCTGTACTGGCACGTGAAGAACAAGCGGGCCCTGCTCGATGCCCTGCCAATCGAGATGCTGGACAGGCATCATACCCACTCCTGCCCCCTGGAAGGCGAGTCATGGCAAGACTTTCTGCGGAACAACGCCAAGTCATACCGCTGTGCTCTCCTCTCACATCGCGACGGGGCTAAAGTGCATCTCGGCACCCGCCCAACAGAGAAACAGTACGAAACCCTGGAAAATCAGCTCGCGTTCCTGTGTCAGCAAGGCTTCTCCCTGGAGAACGCACTGTACGCTCTGTCCGCCGTGGGCCACTTTACACTGGGCTGCGTATTGGAGGAACAGGAGCATCAAGTAGCAAAAGAGGAAAGAGAGACACCTACCACCGATTCTATGCCCCCACTTCTGAAACAAGCAATTGAGCTGTTCGACCGGCAGGGAGCCGAACCTGCCTTCCTTTTCGGCCTGGAACTAATCATATGTGGCCTGGAGAAACAGCTAAAGTGCGAAAGCGGCGGGCCGACCGACGCCCTTGACGATTTTGACTTAGACATGCTCCCAGCCGATGCCCTTGACGACTTTGACCTTGATATGCTGCCTGCTGACGCTCTTGACGATTTTGACCTTGACATGCTCCCCGGGTAA

>TRE3Gp

TTTACTCCCTATCAGTGATAGAGAACGTATGAAGAGTTTACTCCCTATCAGTGATAGAGAACGTATGCAGACTTTACTCCCTATCAGTGATAGAGAACGTATAAGGAGTTTACTCCCTATCAGTGATAGAGAACGTATGACCAGTTTACTCCCTATCAGTGATAGAGAACGTATCTACAGTTTACTCCCTATCAGTGATAGAGAACGTATATCCAGTTTACTCCCTATCAGTGATAGAGAACGTATAAGCTTTAGGCGTGTACGGTGGGCGCCTATAAAAGCAGAGCTCGTTTAGTGAACCGTCAGATCGCCTGGAGCAATTCCACAACACTTTTGTCTTATAC

>Sp_pks1_CRE

GAATCATATAAATGGGATTCAGTGCACAAATTCATTGAAGTACCCTTCTAATTCTTCTTTGACATATACATGTATCTATTTATATTTTATGTGAAAGTTGACAAACCTCCATTGAGACAGGAAAGGAAACAGCGTTTTTCAAAGCTTTCTGGTGGTAGTATAATACTTTCAAGGAGGTTTTTTTTCCGCCCGCAACTCTCATCACTACCGGACGTCAGTGCTGAGTTAAGTTACCCGTAGTTAATAAAAGAAATAACCGACAGTTGATGCTGACAGCAGGGGCCCAGGATATTTCTGGTTTCTACTAATTTGTATAACTTGCATTCCTCCACGCCGGGGGAACGTGTTACTATAATACTCTTATGAGTTATGAGATCCGGCGCCGGAGTGTTTTTAAGGGGTATTCGTCTTTAAAAAAAAAGAGATAATATCTGACAAGTGAGGAACAGCAAGTACAACAACG

>Sp_fmo2_CRE

GCGGATTTGCCCCGGCCTGCCCCCGAGGGCCAATACAAGAAATGGCACGAAGCTCGTAGATAATGCAAACACAAACATCGTGGCCCGTAGCCTCCAATTTGTGCATACTTATAAAAGTAGCTTATCAAACTAATCTTTATGAAAGATTTCGACTCAAGTACAACCTTGTACCAGCATTTTAAAACGGTACTAAATTTACCAGCGAACAGGATAAAGCAGTCGTTAATTGTTACGAGGATATGGCACGAAATACACGTAAGTGCAGTGACGTAGATACAGGGCGCGTACCCCTCTCCCACAACGCTAGAGCGTCCTAGTGGCTGGACTACTCACGGTACACGCATGACGTATGCTAAATAAAAAAAAAATCCATTCGTCGATCTGTTGCAAGTCGAAACAT

>Sp_endo16_CRE

<u>GCTTCGTCGATATCGTCATA</u>TTACTGAAGGGGATAAACATTCTTGCTGCAGTTCTCGAGGTTGCGTAAAGTAGGGACGACCGAGAAACAAAATGTCTGCACAGAATTAAAGTGACAGCAAATGTATGTTCTTAATTTAGGTTAATTAATAATTAGTAGGAAGAAGAAAATGGGCCTTTGCTAAGCAGTGAACAAACAAACGACCAAAGTGGGCAACAAAACAGAATAAAGAAAAACAACAACGAGATTAACGAAAATATAGAAAAAAGGAAAGAAAAGAAGGACAAAGAAGTGTAGATAGAGAGAGGAAAACCAACAAAGAGAGCGCGACAGAGAGAGAGAGAGAGAGAGAAAACCTGGCGAAACGATAGGGATATACATGTATATATAGAGAGAAAGAGAGACAGAGAACAAGAGAGAGAGAGAGAGAGAGAGAGAGAGAGAGAGAGAAAGGGATAGATAGAGATAGAGAGAGAGAGAGAGAGAGATGGGTGTTTCTTATGACATGTTGCTGTACACGTAGTAGGCCACATATTACAGTACTGTCAAAATTGCAAATTTTATGTTTTCGGCATCTTTATCCGACACAATTATCAGCGATTATACCACATACATCCGATGATGCACTCGTCACAACCGGTATTATATACTTCATGCTTGCTTGCTAAGACAGAATGTGTATTTATGCTTGCAAGTACAGTGTAAGCAACCAGGGGACCAACAACTAAAAAGTCCTCTCCGAGGGACTTGGTAATAAAGATTAATGCCTTACCAAAACGCACTAGCTTATCCACTTCCGGGTCCTTCTATTATTACGTGACCCTTGCTCGACCACTTGCTCTCGTCTATCGTCAATAAAGATTATGTACAGATCCCGTTAATGAGTTTACGAACAGTGTCTCTCTCACAATTGTCTCGATATTTGTCCAGATTACATTAAGAACAATGATCACTAAAAGGATTTTTGTCGGTCCTTCATTATTTTTCTTCCTGCTAGTAGGCCTAATTGACATTTATTAGTCATACTTCTCATATTTTATATGTTATCGTGGTTTTATCTTATCCTTCCTGTTCCTGCCCGACCTATAGCAACTTTCTTGTTCATTATTTCAAATCACTGTACCTAGAATTTAAAGATTTGACATTGTTTAGCACTTAATTTGGTAACGAACGTCAATCTTAAGTAATACAGACGAAACTCGATCACAACCTTTTGCTTGACTTTTATATTAACCTTAGAATATTATCGCCAATCTAGATGGTTCTATAGAATTGGGTCGGTGACCTAATTTCCCTTGTTACGCAGTTTTGTATATCGGATATCGTGACATTAATTTATAATATATGATGACATTTTTGCTTCATATTTTGCGGTACCGGGGGATGGTGATTTTAACATGGGGATAAAGATATTGCATCAAGATTTGCACAAGCTCTTGTTCTAATATCCACCTTGCCCCCCCCCCCCCATCTGCTCCCCTCTCACTCCCTATTTCTTCTTCTTCGTCTTCTTCTTCTTCGTCTTCTCCTTTTCCCCTTTGTACATATATCATTTTCTTTATCCTCTCTCTCTCTCCCTCGTTTCCATTTACACCTTCCTCTGTTTCTGTTTCTGTATCTCACTCTTTTTTCTTACACTGACCCAGTTCGCACTCCCTCTTTATTTTTCCTTCACCCAGACTCGATCTCTCTCTCTCTCTCCTGCTCTATATCTCTCTGTATATCTGTCTCTATGTGTGTGTGGGTGTGTGTATGTGTGTGTGCGTGCGTGTGTGTGCGTGTGTGTGCGTGTGTGTGTGTGTGTGTGTCTCTCTCTCTCTCTCTCTTTCTCTCTCTCTCACTCTCTAAGTATATATCTCCTTCCCCATTTTCTCTTTCCCACTTTTCTCTTTCCCCCTCTCAAATATTGATAAAAAGAATACATAATTTGGGTTTTCTGTTGTACGCAGAAAACCTAAATGTCGTATTCCTTCACAAATATTCGACTTCGAACACATTCCTTGCAGAAATGTGTCTCTAATCACATCCTCCTAATACATTATGATACAATTTTATTTAAGGAAAATGTTGTCGTCAAATGTATGGGGCTCCCAACGCTTCAAAGGGGCTTTAAAGTTATCATATGAATGTAACCTAAACCTTCTGAATTACTGTCATGATATTGGGCACTGCTGGGATGATTTTATCAATGACCAAACCGTAACTTTTGATAAAATGTCATTGCGCGTAAAGTAGACGACCGCCCCTCCTCTTCCTTCCTTTCGAGTCGTTGATCCTCCCTCCAAAAAAAGTCTTATTATGACGAAATAATAAGTATGAATAGTATTAGGAACAGATAGTATCTCGATACGGAGTCTTGTTGAATACAATGCTTATACACGATAATGTGGACGCACTTTGCACACTGTTATCTATCAAATTTCTTCAAAGGAACATGTCTAGCCTTGTTTTACCGACATAAACAGCCAGCTTTGGTTGCTAAGTGTAGTGCAGACGGCGGATCGGGTTAAAGCATGAATAAGATATGTTTTAAGTCTTTGGTTTTCTCTAAACTGTGTGTGATAACAGAAGAAAATTGATGTTTTAGTTACAGGAACATATTTTGGAGTAGGTAACAGGTATTGGTATACTGGCTCCTGGTACACTAATGTACACTCACACTCAAAACGTGGCTTATGGAGAAGGGTTTGGATTTCCAATTCGGGGTTGTTTTTGTATATCAAATAACAAATGAAGAGGGCACTTCTAATTCAGAAACGTTATCATGAATAAAGACTTTAACTTTGTTGGTTTTTCAAATTAATTTCGGATGTTTTTCAGTCGATTCCGATGAGATAAAACCCCGAATATACCTGGAACTGAACGTTCTTTGTTTCCACGAAAGATTTAATGCCCGTAAACACAAACATCTGACAAATGCATTAAACTTCATCAAACCATGTAACAAAAGCGTAATTCCTCTTAAATCGCCCTTACTTCTAAGAACGTCGTAAATCGCAACGTCTCAAAAATATTGACCAAAATGATCACGCCATTATCATCGCGGAGGATTAAGTGATCAAGCTACCAAGAGATTACATCATTTCAAAGTTATCGCATCCCAGGTTAAACTGTTTGAGTTTTGGATTGTGCTATCAAAGACAAAGGGGTGTAACTTTACCCCCCTCATCAAGAGCGGAGGGTTAAATAGAGAAAGACTGGTCGAGGACAGGTCATAATATTGCTAATTTTTAAGCTTATCATC

>Sp_nodal_CRE

GGTAATGTTTGGCAGGGGTATTTTATTTAAAGTCTAATTTACTCATGTGTTCTGTCTTTGTTTGGATTAACGAAACATGTTTCTTTTACCAAATTATTATTTCTGCGATTTACCACTTCAGTTTCCTCCCTTTAAACCTGTTCCGGTAATTGACTGACTAACAAACTATGTGGAGAAACATTATTTGAACGGAATTCTCCTTTCCTTTATTTGGATCGTAATGAACCCACTATAGCCCTGATTAAACATCCTTATCTAATTAAAACATTTATAAAAACAAAACTTGTTTGCTGTATTTTCGGAAGTGGATTAGTGCCATTGAAACATAAAAGTACTTCTTTAAACTCGTTTGTTTTTAATTGTGTGCTCAATGTTTGGTTTGGTTTTGTTTGTTTGGTCGTGCGCTTTCCGTATATAGAACAAAGATGTGCCTTTTTTTCGTTCCCTTTTGGTAGAATGAAGATACACTTTCCCTTTTAATAGTGTTTAAGTTAATGTGCACGCAACATGATGGCTTTGCATTTTGCGATATATTGTACAACCATGTATAAGAGATCAATAATGGAATGGAATGGCGGAGGGGGGGGGGGGGGGGAAGAATGTTTTTCTTCTCATAGTGACAAAAATCCTACATCAGTTTCCCCCTCGCTTTCACTCTTCTGCACTTGCTGTTGTGTCTCTTTTTACCCCCTTCTCTCTCACTCTCTCCTTCTATGCCCATTCTCTCTCTCTCTTTCTATCCCACTCTCCTTCTACACCTACTCTCACCCTCCCTATATGCCCCCCACCCTTTCTCTCTCTCTCTCTCTCTCCCTCTCCTTCTCCTTCTCTCTCCCCCATCTCTCTCTATCTCTCTCTCCCTCTCGCCATTTTGTCTCTCCCTGTATCTGTCTTAATCTCTTTACCTTGCCTTCTTCTATATACGTCTCCCTTCTTTCGTTCTCCATTCCTCAGTATGTGTAATATCGCCGCAATCCCCATCCATGAAATACAATTGTTCTAATTTCAGGTTAGTTCCATCATTTTTTCACGAAATAAAGCCCAAGTTCAATATGTAGCTGAAAGCGAGCCGATCTTGTAAACTCGAGTTCTTGTCACGGTGTTCAAAGCAAAGGTGTATTAATTCATTCTGGGAAAATATATTTTGATTGTAATTGCATTTCTTTTTCTCACAGACAATACTATTTTTTTAAACCTAGGGCTTATGTCAATAATGAGAAATATAGACAATCTGGTTTTTATTTTAAAAAGGGCGTTTCCTCAACACCTCTGTATCAGCACAATTTACAAACATGCCGAATGATTTCCACTAAAATGAAACACTCAAGCCTGATATAATATTTACGATTAGCATGCCGATTTATTTACATTCAAACACAAGTGTAAACTAGAGGTAGAATTGCCTTAAAAATACATTTGTTATGATTATCTGCCATCGAGTAAAATTTATTGAAGAAATAAATTGATAAAGCAGTTTTATTTGGTAACAAATTACTAAATTGTCGATTCATGAGTCTCTATTTCTTTTACCGCCATCAAAACAAAATAATTGCAAAGATTCCATATTATTTGCACAGTAGCAAATATTCATTATTCTATATCTATGTTGTATTAATATATTTGATCAGATGATTATTGCATGCTCAAAACATGAATAAATTTCATATGTGCAAAAATCGTATATTATCGTATTTCCGAAAATTCCCGATGAATAATTATGCAAATATATTCACATTATTTTTGTAACTTCTTATTTTCATTTCTTTTCTGTTGTTAAACGAGTGTATGAAGCCCACATTCTGACGAGACTGTTGTGCTTTCCTTAGATCACAACACATGAAGCCAATTGTTTTCAAAGCTGCCAATTAAAGCAGTCTGAAGCTCAGCTAAACATTATTCACCGAATTGTACTCTTCAAAATGAAGTATCGAGAAAGAAGATAGACTAATGCGTAACACTTAGCAGGACTACCCTTTTATTGTCGTTTGGTTACAACTAATTATAGTGTGCTTAGAGTCCTACAAAAGAGCCGATTCATTGTTAATTAGGAAGACGGGATGGGTACATGTCGAGCAGCCGGGATGGGTTCTATGCATAGTGGCCCGAGAACCGTTGCTTTCATTCACCGTGGTTCGGCTGAATGGTTTTTAGCATGAATGCTTGAATTGCGAAACAATGCGCGCGCATGTTGAGCTAGGGGTCTTTGAAAAGAGTGTCTGATTAGTGTGGCAAGGCGAGAGTAGTGTCTGGGAAGGTAAGGTCTCAAGTATTTAAGATGTCTGGCTCACTTTCACCTTCATTATTTCGTCAGAAACTCTAACCAAGACAACATCACGCTAACAATTTCTCAAGTACTCTCAAATTTGAAGTAAAAAATTAATTAAAGTCACAAAGTGTGTTTGTGCAAGCTTCGACAAGAGATTATTGAAAGATTACCTGCAACGCTCATCATCAGCCTGGACTCACCTCATCATCAGCCTGGACTCATCTCATCAAAGGGTTTCTACATTTCAGAGAGCTGAGACATCCCAGTGACGACATCGTTCCAGCGAAGCTTCGGAATATTTTTAAGGACTTTTAACC

### Primers for generating templates for in vitro transcription reaction

#### DmEn.GFP or dnLv-Ets1.GFP mRNA

>Sp6_GFP_Forward

TATAATTTAGGTGACACTATAGAAGAG<u>ACTAACATACGCTCTCCATC</u>

#### Sp-caMEK mRNA

>SP6_SpMEK_Forward

TATAATTTAGGTGACACTATAGAAGAGTATATATATCATCATGTCGAAGACGCCCGGAAA

#### Sp-DUSP6 mRNA

>SP6_SpDUSP6_Forward

TATAATTTAGGTGACACTATAGAAGAGTATATATATCATCATGCCCGCTCAGACTCCTGA

#### Universal reverse primer

>Sp6_Reverse

CGGAAGGAGCTGACTGGGTT

### RNA probe primers p16

>p16_F

CCATCATCGCTTTATTTGC

>p16_T3_R

<u>AATTAACCCTCACTAAAGGGAGA</u>CTGTTGTCTTCTGACGAGTC

### sm30b

>sm30b_F

GCCTGGCAGAGAACAACTAT

>sm30b_T3_R

<u>AATTAACCCTCACTAAAGGGAGA</u>GTCCATCCCAAGCGTAAAGA

## Notes

### Competing Interest Statement

The authors have declared no competing interest.

